# Learning optimal decisions with confidence

**DOI:** 10.1101/244269

**Authors:** Jan Drugowitsch, André G. Mendonça, Zachary F. Mainen, Alexandre Pouget

**Author notes:** Corresponding author: Jan Drugowitsch, Department of Neurobiology, Harvard Medical School, 220 Longwood Avenue, Boston, MA 02115.

## Abstract

Diffusion decision models (DDMs) are immensely successful models for decision-making under uncertainty and time pressure. In the context of perceptual decision making, these models typically start with two input units, organized in a neuron-antineuron pair. In contrast, in the brain, sensory inputs are encoded through the activity of large neuronal populations. Moreover, while DDMs are wired by hand, the nervous system must learn the weights of the network through trial and error. There is currently no normative theory of learning in DDMs and therefore no theory of how decision makers could learn to make optimal decisions in this context. Here, we derive the first such rule for learning a near-optimal linear combination of DDM inputs based on trial-by-trial feedback. The rule is Bayesian in the sense that it learns not only the mean of the weights but also the uncertainty around this mean in the form of a covariance matrix. In this rule, the rate of learning is proportional (resp. inversely proportional) to confidence for incorrect (resp. correct) decisions. Furthermore, we show that, in volatile environments, the rule predicts a bias towards repeating the same choice after correct decisions, with a bias strength that is modulated by the previous choice’s difficulty. Finally, we extend our learning rule to cases for which one of the choices is more likely a priori, which provides new insights into how such biases modulate the mechanisms leading to optimal decisions in diffusion models.

**Significance Statement:** Popular models for the tradeoff between speed and accuracy of everyday decisions usually assume fixed, low-dimensional sensory inputs. In contrast, in the brain, these inputs are distributed across larger populations of neurons, and their interpretation needs to be learned from feedback. We ask how such learning could occur and demonstrate that efficient learning is significantly modulated by decision confidence. This modulation predicts a particular dependency pattern between consecutive choices, and provides new insight into how a priori biases for particular choices modulate the mechanisms leading to efficient decisions in these models.

## Introduction

Decisions are a ubiquitous component of every-day behavior. To be efficient, they require handling the uncertainty arising from the noisy and ambiguous information that the environment provides (1). This is reflected in the trade-off between speed and accuracy of decisions. Fast choices rely on little information and may therefore sacrifice accuracy. In contrast, slow choices provide more opportunity to accumulate evidence and thus may be more likely to be correct, but are more costly in terms of attention or effort and lost time and opportunity. Therefore, efficient decisions require not only a mechanism to accumulate evidence, but also one to trigger a choice once enough evidence has been collected. *Drift-diffusion models* (or *diffusion decision models*; DDMs) are a widely-used model family (2) that provides both mechanisms. Not only do DDMs yield surprisingly good fits to human and animal behavior (3–5), but they are also known to achieve a Bayes-optimal decision strategy under a wide range of circumstances (4, 6–10).

DDMs assume a particle that drifts and diffuses until it reaches one of two boundaries, each triggering a different choice (Fig. 1a). The particle’s drift reflects the net surplus of evidence towards one of two choices. This is exemplified by the random-dot motion task, in which the motion direction and coherence set the drift sign and magnitude. The particle’s stochastic diffusion reflects the uncertainty in the momentary evidence and is responsible for the variability in decision times and choices widely observed in human and animal decisions (3, 5). A standard assumption underlying DDMs is that the noisy momentary evidence that is accumulated over time is one-dimensional — an abstraction of the momentary decision-related evidence of some stimulus. In reality, however, evidence would usually be distributed across a larger number of inputs, such as a neural population in the brain, rather than individual neurons (or neuron/anti-neuron pairs; Fig. 1a). Furthermore, the brain would not know a priori how this distributed encoding provides information about the correctness of either choice. As a consequence, it needs to learn how to interpret neural population activity from the success and failure of previous choices. How such an interpretation can be efficiently learned over time, both normatively and mechanistically, is the focus of this work.

**Figure 1.**
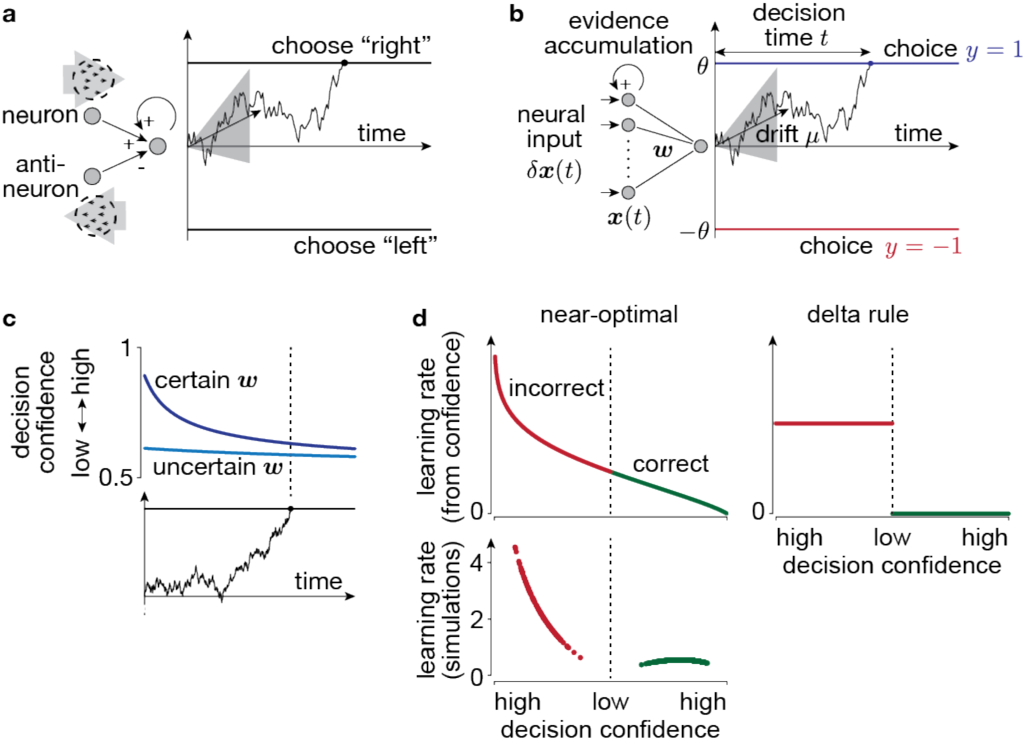
Learning the input weights from feedback in diffusion models. In diffusion models, the input(s) provide at each point in time noisy evidence about the world’s true state, here given by the drift *μ*. The decision maker accumulates this evidence over time (e.g., black example traces) to form a belief about *μ*. Bayes-optimal decisions choose according to the sign of the accumulated evidence, justifying the two decision boundaries that trigger opposing choices. (**a**) In standard diffusion models, the momentary evidence either arises directly from noisy samples of *μ*, or, as illustrated here, from a neuron/anti-neuron pair that codes for opposing directions of evidence. The illustrated example assumes a random dot task, in which the decision maker needs to identify if most of the dots that compose the stimulus are moving either to the left or to the right. The two neurons (or neural pools) are assumed to extract motion energy of this stimulus towards the right (top) and left (bottom), such that their difference forms the momentary evidence towards rightward motion. A decision is made once the accumulated momentary evidence reaches one of two decision boundaries, triggering opposing choices. (**b**) Our setup differs from that in (**a**) in that we assume the input information *δ****x***(*t*) to be encoded in a larger neural population whose activity is linearly combined with weights ***w*** to yield the one-dimensional momentary evidence, and that the decision maker aims to learn these weights from feedback about the correctness of her choices. (**c**) Decision confidence (i.e., the belief that the made choice was correct) in this kind of diffusion model drop as a function of time (horizontal axis) and with increased uncertainty about the input weights (different shades of blue). (**d**) For near-optimal learning, the learning rate (the term *ξ*_*w*_ in Eq. (6)) is modulated by decision confidence (top left). High-confidence decisions lead to little learning if correct (green, right), and strong learning if incorrect (red, left). Low-confidence decisions result in a moderate confidence-related learning rate term (top, center). The learning rate in 1000 simulated trials (bottom) shows that the overall learning rate preserves this trend, with an additional suppression of learning for low-confidence decisions. Other learning heuristics (e.g., the delta rule, right) do not modulate their learning by confidence.

The multiple existing computational models for how humans and animals might learn to improve their decisions from feedback (e.g., 11–14) do not address the question we are asking, as they all assume that all evidence for each choice is provided at once, without considering the temporal aspect of evidence accumulation. This is akin to fixed-duration experiments, in which the evidence accumulation time is determined by the environment rather than the decision maker. We, instead, address a more general and natural case in which decision times are under the decision maker’s control. In this setting, commonly studied using “reaction time” paradigms, the temporal accumulation of evidence needs to be treated explicitly, and – as we will show – the time it took to accumulate this evidence impacts how the decision strategy is updated after feedback. Some models for both choice and reaction times have addressed the presence of high-dimensional inputs (e.g., 15–17). However, they usually assumed as many choices as inputs, were mechanistic rather than normative, and did not consider how interpreting the input could be learned. We furthermore extend on previous work by considering the effect of a priori biases towards believing that one option is more correct than the other, and how such biases can be learned. This yields a new theoretical understanding of how choice biases impact optimal decision-making in diffusion models. Furthermore, it clarifies of how different implementations of this bias result in different diffusion model implementations, like the one proposed by Hanks et al. (18).

## Results

### Bayes-optimal decision-making with diffusion models

A standard way (8, 10, 19) to interpret diffusion models as mechanistic implementations of Bayes-optimal decision-making is to assume that, in each trial, an unobservable latent state *μ* (called *drift rate* in diffusion models) is drawn from a prior distribution, 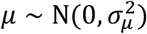, with zero mean and variance 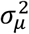. The decision maker’s aim is to infer whether this latent state is positive or negative (e.g., rightward vs. leftward motion in the random dot motion task), irrespective of its magnitude (e.g., the dot coherence level). The latent state itself is not directly observed, but is indirectly conveyed via a stream of noisy, momentary evidence values *δz*_1_, *δz*_2_, …, that, in each small time step of size *δt*, provide independent and identically distributed noisy information about *μ* through *δz*_*i*_|*μ* ∼ N(*μδt*, *δt*). Here, we have chosen a unit variance, scaled by *δt*. Any re-scaling of this variance by an additional parameter would result in a global re-scaling of the evidence that can be factored out (4, 8, 20), thus making such a re-scaling unnecessary.

Having after some time *t* ≡ *nδt* observed *n* pieces of such evidence, *δz*_1:*n*_, the decision maker’s posterior belief about *μ*, *p*(*μ*|*δz*_1:*n*_), turns out to be fully determined by the accumulated evidence, 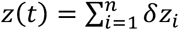, and time *t* (see Methods). Then, the posterior belief about *μ* being positive (e.g., leftward motion) results in (8)

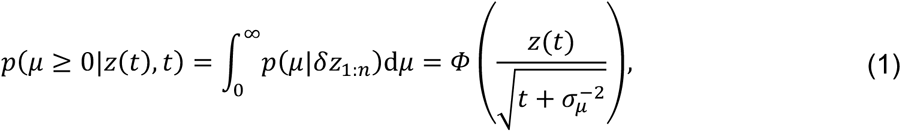

where *Φ*(⋅) is the cumulative function of a standard Gaussian. The opposite belief about *μ* being negative is simply *p*(*μ* < 0|*z*(*t*), *t*) = 1 − *p*(*μ* ≥ 0|*z*(*t*), *t*) (Fig. 3a). The accumulated evidence follows a diffusion process, *z*(*t*)|*μ* ∼ *N*(*μt*, *t*), and thus can be interpreted as the location of a drifting and diffusing particle with drift *μ* and unit diffusion variance (Fig. 1a). By Eq. (1), the posterior belief about *μ* ≥ 0 is > 1/2 for positive *z*(*t*), and < 1/2 for negative *z*(*t*). To make Bayes-optimal decisions, Bayesian decision theory (21) requires that these decisions are chosen to maximize the expected associated reward (or, more formally, to minimize the expected loss). Assuming equally-rewarding correct choices, this implies choosing the option that is considered more likely correct. Given the above posterior belief, this makes *y* = sign(*z*(*t*)) ∈ {−1,1} the Bayes-optimal choice, which can be implemented mechanistically by (possibly time-varying) boundaries ±*θ*(*t*) on *z*(*t*), associated with the two choices. At these boundaries, the posterior belief about having made the correct choice, or decision confidence (22), is then given by Eq. (1) with *z*(*t*) replaced by *θ*(*t*). The sufficient statistics, *z*(*t*) and *t*, of this posterior remain unchanged by the introduction of such decision boundaries, such that Eq. (1) remains valid even in the presence of these boundaries (8). Thus, under the above assumptions of prior and evidence, diffusion models implement the Bayes-optimal decision strategy (Fig. 3b).

Note that |*μ*| (i.e., the momentary evidence’s signal-to-noise ratio) controls the amount of information provided about the sign of *μ*, and thus the difficulty of individual decisions. Thus, the used prior 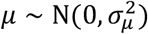, which has more mass on small |*μ*|, reflects that the difficulty of decisions varies across trials, and that harder decisions are more frequent than easier ones. The prior width, 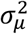 determines the spread of *μ*’s across trials, and therefore the overall difficulty of the task (larger 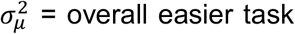). We chose a Gaussian prior for mathematical convenience, and also because hard trials are more frequent than easy ones in many experiments (e.g. (20)), even though they don’t commonly use Gaussian priors. In general, the important assumption is that the difficulty varies across trials, but not exactly how it does so, which is to say that the shape of the prior distribution is not critical (8). Different prior choice will not qualitatively change our results, but would make it hard or impossible to derive interpretable closed-form expressions. Model predictions would change qualitatively if we assume the difficulty to be fixed, or known a-priori (see 8), but we will not consider this case, as it rarely if ever occurs in the real world.

### Using high-dimensional diffusion model inputs

To extend diffusion models to multi-dimensional momentary evidence, we assume it to be given the a *k*-dimensional vector *δ****x***_*i*_. This evidence might represent inputs from multiple sensors, or the (abstract) activity of a neuronal population (Fig. 1b). As the activity of neurons in a population that encodes limited information about the latent state *μ* is likely correlated across neurons (23, 24), we chose the momentary evidence statistics to also feature such correlations (see Methods). In general, we choose these statistics such that ***w***^*T*^*δ****x***_4_ = *δz*_*i*_, where the vector ***w*** denote the *k* input weights (for now assumed known). Defining the high-dimensional accumulated evidence by 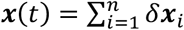, this implies *z*(*t*) = ***w***^*T*^***x***(*t*), such that it is again Bayes-optimal to trigger decisions as soon as ***w***^*T*^***x***(*t*) equals one of two decision boundaries ±*θ*(*t*). Furthermore, the posterior belief about *μ* ≥ 0 is, similar to Eq. (1), given by

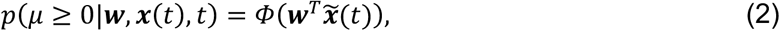

where we have defined the time-attenuated accumulated evidence 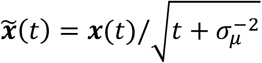. As a consequence, the decision-confidence for either choice, is, as before, given by Eq. (2), with 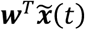 replaced by 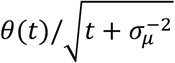. For time-independent decision bounds, *θ*(*t*) = *θ*, this confidence decreases over time (Fig. 1c), reflecting the uncertainty about *μ*, and that late choices are likely due to a low *μ*, which is associated with a hard trial, and thus low decision confidence. This counter-intuitive drop in confidence with time has been previously described for diffusion models with one-dimensional inputs (8, 25), and is a consequence of a trial difficulty that varies across trials. Specifically, it arises from a mixture of easy trials associated with large |*μ*| that lead to rapid, high-confidence choice, and hard trials associated with small |*μ*| that lead to slow, low-confidence choices. Therefore, it doesn’t depend on our choice of Gaussian prior, but is present for any choice of symmetric prior over *μ* (see SI). The confidence remains constant over time only when the difficulty is fixed across trials (i.e., *μ* ∈ {−*μ*_0_, *μ*_0_} for some fixed *μ*_0_).

### Using feedback to find the posterior weights

So far we have assumed the decision maker knows the linear input weights ***w*** to make Bayes-optimal choices. If they were not known, how could they be learned? Traditionally, learning has been considered an optimization problem, in which the decision maker tunes some decision-making parameters (here, the input weights ***w***) to maximize their performance. Here we will instead consider it as an inference problem in which the decision maker aims to identify the decision-making parameters that are most compatible with the provided observations. These two views are not necessarily incompatible. For example, minimizing the mean squared error of a linear model (an optimization problem) yields the same solution as sequential Bayesian linear regression (an inference problem) (26). In fact, as we show in the SI, our learning problem can also be formulated as an optimization problem. Nonetheless, we here take the learning-by-inference route, as it provides a statistical interpretation of the involved quantities, which provides additional insights. Specifically, we focus on learning the weights while keeping the diffusion model boundaries fixed. The decision maker’s reward rate (i.e., average number of correct choices per unit time), which we use as our performance measure, depends on both weights and the chosen decision boundaries. However, to isolate the problem of weight learning, we fix the boundaries such that a particular set of optimal weights ***w***^∗^ maximize this reward rate. The aim of weight learning is to find these weights. Weight learning is a problem that needs to be solved even if the decision boundaries are optimized at the same time. We have addressed how to best tune these boundaries elsewhere (8, 27).

To see how learning can be treated as inference, consider the following scenario. Before having observed any evidence, the decision maker has some belief, *p*(***w***), about the input weights, either as a prior or formed through previous experience. They now observe new evidence, *δ****x***_1_, *δ****x***_2_, … and use the mean of the belief over weights, 〈***w***〉 (or any other statistics), to combine this evidence and to trigger a choice *y* once the combined evidence reaches one of the decision boundaries. Upon this choice, they receive feedback *y*^∗^ about which choice was the correct one. Then, the best way to update the belief about ***w*** in light of this feedback is by Bayes’ rule,

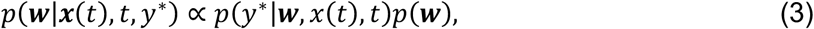

where we have replaced the stream of evidence *δ****x***_1_, *δ****x***_2_, … by the previously established sufficient statistics ***x***(*t*) and *t*.

The likelihood *p*(*y*^∗^|***w***, ***x***(*t*), *t*) expresses for any hypothetical weight vector ***w*** the probability that the observed evidence makes *y*^∗^ the correct choice. To find its functional form, consider that, for a known weight vector, we have shown that *p*(*μ* ≥ 0|***w***, ***x***(*t*), *t*), given by Eq. (2), expresses the probability that *y* = 1 (associated with *μ* ≥ 0) is the correct choice. Therefore, 1 − *p*(*μ* ≥ 0|***w***, ***x***(*t*), *t*) corresponds to the probability that *y* = −1 (associated with *μ* < 0) is the correct choice. Therefore, it can act as the above likelihood function, which, by Eq.(2), is given by 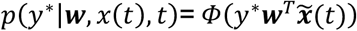, where we have used 1 − *Φ*(*a*) = *Φ*(−*a*). In summary, the decision maker’s belief is optimally updated after each choice by

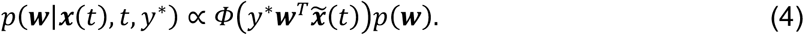

This update equation only requires knowing the accumulated evidence ***x***(*t*), decision time *t*, and feedback *y*^∗^, but is independent of the chosen option *y*, and how the decision maker came to this choice. As a matter of fact, the decision maker could make random choices, irrespective of the accumulated evidence, and still learn ***w*** according to the above update equation, as long as they keep track of ***x***(*t*) and *t*, and acknowledge the feedback *y*^∗^. Therefore, learning and decision-making aren’t necessarily coupled. Nonetheless, we assume for all simulations that decision makers perform decisions by using the mean estimate 〈***w***〉, which is an intuitively sensible choice if the decision maker’s aim is to maximize their reward rate (see SI).

As in Eq. (4) the likelihood parameters, ***w***, are linear within a cumulative Gaussian function, such problems are known as *Probit Regression* and don’t have a closed-form expression for the posterior. We could proceed by sampling from the posterior by Markov Chain Monte Carlo methods, but that would not provide much insight into the different factors that modulate learning the posterior weights. Instead, we proceed by deriving a closed-form approximation to this posterior to provide such insight, as well as a potential mechanistic implementation.

### Confidence controls the learning rate

To find an approximation to the posterior in Eq. (4), let us assume the prior to be given by the Gaussian distribution, *p*(***w***) = *N*(***w***|***μ***_*w*_, ***Σ***_*w*_), with mean ***μ***_*w*_ and covariance **Σ**_*w*_, which is the maximum entropy distribution that specifies the mean and covariance (28). First, we investigated how knowing ***w*** with limited certainty, as specified by **Σ**_*w*_, impacts the decision confidence. Marginalizing over all possible ***w***’s (see Methods) resulted in the choice confidence to be given by

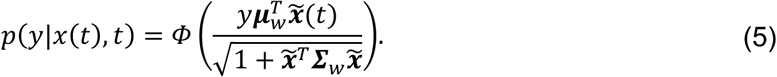

Compared to Eq. (2), the choice confidence is additionally attenuated by **Σ**_*w*_. Specifically, higher weight uncertainty (i.e., an overall larger covariance **Σ**_*w*_) results in a lower decision confidence, as one would intuitively expect (Fig. 1c).

Next, we found a closed-form approximation to the posterior, Eq. (4). For repeated learning across consecutive decisions, the posterior over the weights after the previous decision becomes the prior for the new decision. Unfortunately, a direct application of this principle would lead to a posterior that changes its functional form after each update, making it intractable. We instead used *Assumed Density Filtering* (ADF) (26, 29) that posits a fixed functional form 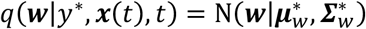 of the posterior density – in our case Gaussian for consistency with the prior – and then finds the posterior parameters 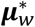 and 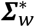 that make this approximate posterior best match the “true” posterior *p*(***w***|*y*^∗^, ***x***(*t*), *t*), Eq. (4). Performing this match by minimizing the Kullback-Leiber divergence KL(*p*|*q*) results in the posterior mean (30, 31)

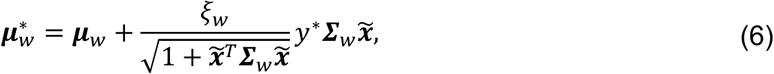

and a similar expression for the posterior covariance (see Methods). Choosing *KL*(*p*|*q*) to measure the distance between *p* and *q* is to some degree arbitrary, but has beneficial properties, such as that it causes the first two moments of *q* to match those of *p* (see SI). In Eq. (6), the factor *ξ*_*w*_ modulates how strongly this mean is updated towards 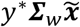, and turns out to be a monotonically decreasing function of decision confidence (Fig. 1d, top; see Methods for mathematical expression). For incorrect choices, for which the decision confidence is *p*(*y*^∗|^***x***(*t*), *t*) < 1/2, *ξ*_*w*_ is largest for choices made with high confidence, promoting significant weight adjustments. For low-confidence choices it only promotes moderate adjustments, notably irrespective of whether the choice was correct or incorrect. High-confidence, correct choices yield a low *ξ*_*w*_, and thus an intuitively minor strategy update. The update of the posterior covariance follows a similar confidence-weighted learning rate modulation (**Fig. S1**; Methods).

Decision confidence is not the only factor that impacts the learning rate in Eq. (6). For instance, 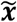 shrinks for longer, less confidence choices (because it is inversely proportional to time) and results in overall less learning. Less certain weights, associated with larger magnitudes of **Σ**_*w*_, have a similar effect. To investigate the overall impact of all of these factors combined on the learning rate, we simulated a long sequence of consecutive choices and plotted the learning rate for a random subset of these trials against the decision confidence (Fig. 1d, bottom). This plot revealed a slight down-weighting of the learning rate for low-confidence choices when compared to *ξ*_*w*_, but left the overall dependency on *ξ*_*w*_ otherwise unchanged.

### Performance comparison to optimal inference and to simpler heuristics

The intuitions provided by near-optimal ADF learning are only informative if its approximations do not cause a significant performance drop. We quantified this drop by comparing ADF performance to that of the Bayes-optimal rule, as found by Gibbs sampling (see Methods). Gibbs sampling is biologically implausible as it requires a complete memory of inputs and feedbacks for past decisions and is intractable for longer decision sequences, but nonetheless provides an optimal baseline to compare against. We furthermore tested the performance of two additional approximations. One was an ADF variant that assumes a diagonal covariance matrix **Σ**_*w*_, yielding a local learning rule that could be implemented by the nervous system. This variant furthermore reduced the number of parameters from quadratic to linear in the size of ***w***. The second was a second-order Taylor expansion of the log-posterior, resulting in a learning rule similar to ADF, but with a lower impact of weight uncertainty on the learning rate (see Methods).

Furthermore, we tested if simpler learning heuristics can match ADF performance. We focused on three rules of increasing complexity. The *delta rule*, which can be considered a variant of temporal-difference learning, or reinforcement learning (32), updates its weight estimate after the *n*th decision by

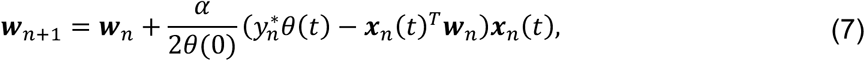

where 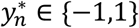 is the feedback about the correct choice provided after this decision, and we have chosen to normalize the learning rate *α* by the initial bound height *θ*(0) to make it less sensitive to this chosen height. As decisions are triggered at one of the two boundaries, ***x***_*n*_(*t*^*T*^***w***_*n*_ ∈ {−*θ*(*t*), *θ*(*t*)}, the residual in brackets is zero for correct choices, and ±2*θ*(*t*) for incorrect choices. As a result, and in contrast to ADF, weight adjustments are only performed after incorrect choices, and with a fixed learning rate *α* rather than one modulated by confidence (Fig. 1d; right). Our simulations revealed that the delta rule excessively and suboptimally decrease in the weight size ‖***w***‖ over time, leading to unrealistically long reaction times and equally unrealistic near-zero weights. To counteract this problem, we designed a *normalized delta rule*, that updates the weight estimates as the delta rule, but thereafter normalizes them by ***w*** ← ***w*** ‖***w***∗‖/‖***w***‖ to ensure that its size matches that of the true weights ***w***^∗^. Access to these true weights, ***w***^∗^, makes it an omniscient learning rule that can’t be implemented by a decision maker in practice. Lastly, we tested a learning rule that performs stochastic gradient ascent on the feedback log-likelihood,

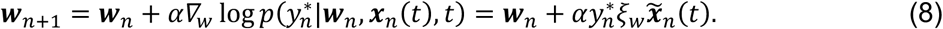

This rule introduces decision confidence weighting through *ξ*_*w*_, but differs from ADF in that it does not take the weight uncertainty (***Σ***_*w*_ in ADF) into account, and requires tuning of the learning rate parameter *α*.

We evaluated the performance of these learning rules by simulating weight learning across 1,000 consecutive decisions (called *trials*; see Methods for details) in a task in which use of the optimal weight vector maximizes the reward rate. This reward rate was the average reward for correct choices minus some small cost for accumulating evidence over the average time across consecutive trials and is a measure we would expect rational decision makers to optimize. For each learning rule we found its reward rate relative to random behavior and optimal choices.

Figure 2a shows this relative reward rate for all learning rules and different numbers of inputs. As can be seen, the performance of ADF and the other probabilistic learning rules is indistinguishable from Bayes-optimal weight learning for all tested numbers of inputs. Surprisingly, the ADF variant that ignores off-diagonal covariance entries even outperformed Bayes-optimal learning for a large number of inputs (Fig. 2a, yellow line for 50 inputs). That reason that a simpler learning rule could outperform the rule deemed optimal by Bayesian decision theory is that this simpler rule has less parameters and a simpler underlying model that was nonetheless good enough to learn the required weights. Learning fewer parameters with the same data resulted in initially better parameter estimates, and better associated performance. Conceptually, this is similar to a linear model outperforming a quadratic model when fitting a quadratic function if little data is available, and if the function is sufficiently close to linear (as illustrated in Fig. S2). Once more data is available, the quadratic model will outperform the linear one. Similarly, the Bayes-optimal learning rule will outperform the simpler one once more feedback has been observed. In our simulation, however, this does not occur within the 1000 simulated trials.

**Figure 2.**
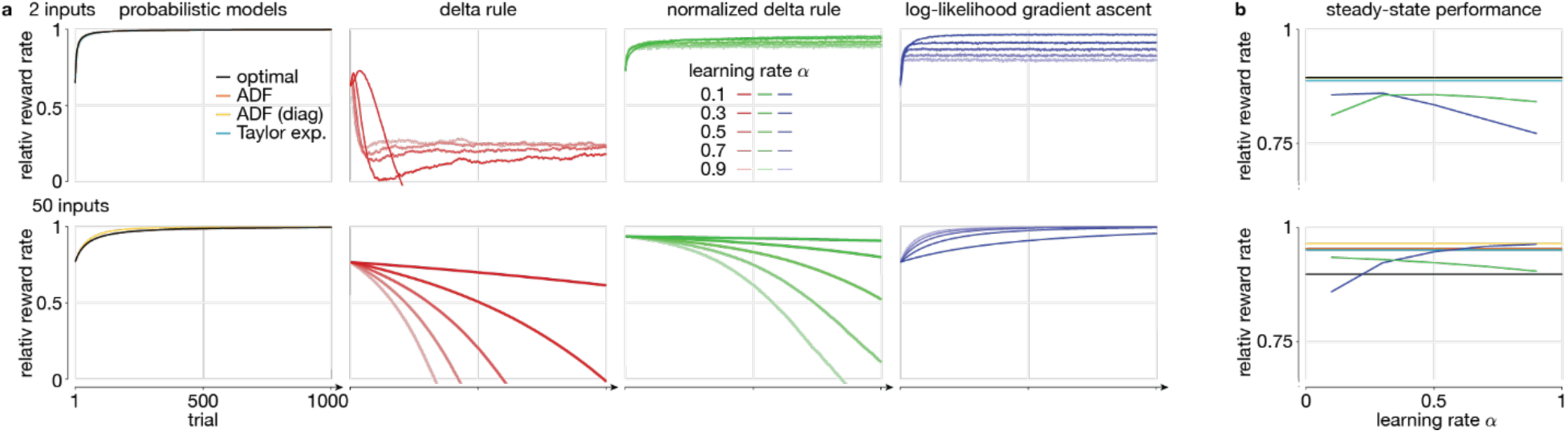
Input weight learning and tracking performance of different learning rules. All plots show the relative reward rate (0 = immediate, random choices, 1 = optimal) averaged over 5,000 simulations with different true, underlying weights, and for 2 (top) and 50 (bottom) inputs. (**a**) The relative reward rate for probabilistic and heuristic learning rules. The probabilistic learning rules include the optimal rule (Gibbs sampling), assumed density filtering (ADF), ADF with a diagonal covariance matrix (ADF (diag)), and a learning rule based on a second-order Taylor expansion of the log-posterior (Taylor exp.). For both 2 and 50 inputs, all rules perform roughly equally. For the heuristic rules, different color shadings indicate different learning rates. The initial performance shown is that *after* the first application of the learning rule, such that initial performances can differ across learning rule. (**b**) The steady-state performance across different heuristic rule learning rates. Steady state performance was measured as an average across 5,000 simulations, averaging over the last 100 of 1000 simulated trials in which the true weights slowly change across consecutive trials. An optimal relative reward rate of one corresponds to knowing the true weight in each trial, which, due to the changing weight, is not achievable in this setup. The color scheme is the same as in (**a**), but the vertical axis has a different scale. The delta rule did not converge and was not included in (**b**).

All other learning heuristics performed significantly worse. For low-dimensional input, the delta rule initially improved its reward rate but worsens it again at a later stage across all learning rates. The normalized delta rule avoided such performance drops for low-dimensional input, but both delta rule variants were unable to cope with high-dimensional inputs. Only stochastic gradient ascent on the log-likelihood provided a stable learning heuristic for high dimensional inputs, but with the downside of having to choose a learning rate. Small learning rates lead to slow learning, and an associated slower drop in angular error. Overall, the probabilistic learning rules significantly outperformed all tested heuristic learning rules and matched (and in one case even exceeded) the weight learning performance of the Bayes-optimal estimator.

### Tracking non-stationary input weights

So far, we have tested how well our weight learning rule is able to learn the true, underlying weights from binary feedback about the correctness of the decision maker’s choices. For this we assumed that the true weights remained constant across decisions. What would happen if these weights change slowly over time? Such a scenario could occur if, for example, the world around us changes slowly, or if the neural representation of this world changes slowly through neural plasticity or similar. In this case, the true weights would become a moving target that we would never be able to learn perfectly. Instead, we would after some initial transient expect to reach steady-state performance that remains roughly constant across consecutive decisions. We compared this steady-state performance of Bayes-optimal learning (now implemented by a particle filter) to that of the probabilistic and heuristic learning rules introduced in the previous section. The probabilistic rules were updated to take into account such a trial-by-trial weight change, as modeled by a first-order autoregressive process (see Methods). The heuristic rules remained unmodified, as their use of a constant learning rate already encapsulates the assumption that the true weights change across decisions.

Figure 2b illustrates the performance of the different learning rules. First, it shows that, for low-dimensional inputs the various probabilistic models yield comparable performances, but for high-dimensional inputs the approximate probabilistic learning rules outperform Bayes-optimal learning. In case of the latter, these approximations weren’t actually harmful, but instead beneficial, for the same reason discussed further above. In particular, the more neurally-realistic ADF variant that only tracked the diagonal of the covariance matrix again outperformed all other probabilistic models. Second, only the heuristic learning rule that performed gradient ascent on the log-likelihood achieved steady-state performance comparable to the approximate probabilistic rules, and then only for high input dimensionality and a specific choice of learning rate. This should come as no surprise, as its use of the likelihood function introduces more task structure information than the other heuristics use. The delta rule did not converge and therefore never achieved steady-state performance. Overall, the ADF variant that focused only on the diagonal covariance matrix achieved the best overall performance.

### Learning both weights and a latent state prior bias

Our learning rule can be generalized to learn prior biases in addition to the input weights. The prior we have used so far for the latent variable, 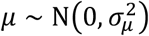, is unbiased, as both *μ* ≥ 0 and *μ* < 0 are equally likely. To introduce a prior bias, we instead used 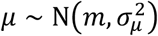, where *m* controls the bias through *P*^+^ ≡ *p*(*μ* ≥ 0) = *Φ*(*m*/*σ*_*μ*_). A positive (or negative) *m* causes *P*^+^ > 1/2 (or < 1/2), thus making *y* = 1 (or *y* = −1) the more likely correct choice even before evidence is accumulated. After evidence accumulation, such a prior results in the posterior

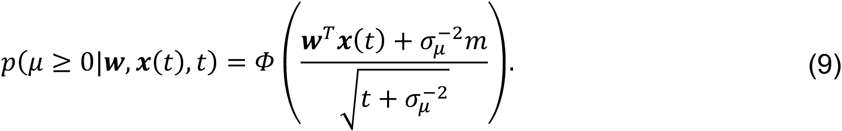

Comparing this to the unbiased posterior Eq. (2) reveals the additional term 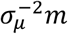 whose relative influence wanes over time.

This additional term has two consequences. First, appending the elements *m* and 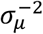 to the vectors ***w*** and ***x***(*t*), respectively, shows that ***w*** and *m* can be learned jointly by the same learning rule we have derived before (Methods). Second, the term requires us to rethink the association between decision boundaries and choices. As Fig. 3c illustrates, such a prior causes a time-invariant shift in the association between the accumulated evidence, *z*(*t*) = ***w***^T^***x***(*t*), and the posterior belief of *μ* ≥ 0 and corresponding decision confidence. This shift makes it possible to have the same Bayes-optimal choice at both decision boundaries (Fig. 3c, blue/red decision areas). Hence, we have lost the mechanistically convenient unique association between decision boundaries and choices. We recover this association by a boundary counter-shift, such that these boundaries come to lie at the same decision confidence levels for opposite choices, making them asymmetric around *z* = 0. Mathematically, this is equivalent to shifting the evidence accumulation starting point, 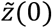 away from zero in the opposite direction (Fig. 3c, shift by 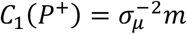; SI). Therefore, a prior bias is implemented by a bias-dependent simple shift of the accumulation starting point, leading to a mechanistically straight-forward implementation of Bayes-optimal decision-making with biased priors.

**Figure 3.**
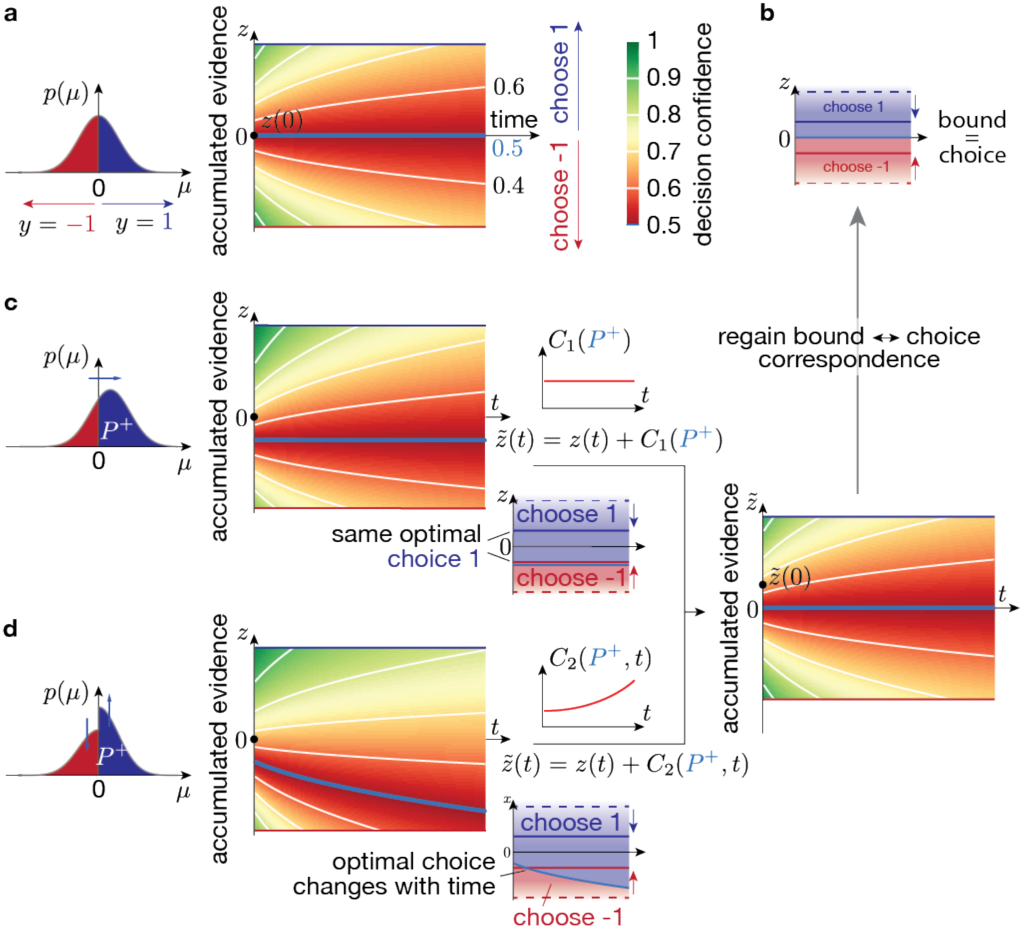
Decision confidence, prior biases, and the relation between decision boundary and choice. (**a**) For an unbiased prior (i.e., *P*^+^ ≡ *p*(*μ* ≥ 0) = 1/2), the decision confidence (color gradient) is symmetric around *z* = 0 for each fixed time *t*. The associated posterior belief *p*(*μ* ≥ 0|*z*(*t*), *t*) (numbers above/below “time” axis label; constant along white lines; ½ along light blue line) promote choosing *y* = 1 and *y* = −1 above (blue area in (**b**)) and below (red area in (**b**)) *z* = 0. (**b**) As a result, different choices are Bayes-optimal at the blue/red decision boundaries, as long as they are separated by *z* = 0, irrespective of the boundary separation (solid vs. dashed blue red lines). (**c**) If the prior is biased by an overall shift, the decision confidence is counter-shifted by the same constant across all *t*. In this case, both decision boundaries might promote the same choice, which can be counter-acted by a time-invariant shift of *z* by *C*_2_(*P*^+^). (**d**) If the prior is biased by boosting one side while suppressing the other, the decision confidence shift becomes time-dependent, such that the optimal choice at a time-invariant boundary might change over time. Counteracting this effect requires a time-dependent shift of *z* by *C*_0_(*P*^+^, *t*). In both (**c**) and (**b**) we have chosen *P*^+^ = 0.6, for illustration.

A consequence of the shifted accumulation starting point is that, for some fixed decision time *t*, the decision confidence at both boundaries is the same (Fig. 3c right). This seems at odds with the intuition that a biased prior ought to bias the decision confidence in favor of the more likely option. However, this mechanism does end up assigning higher average confidence to the more likely option because of reactions times. As the starting point is now further away from the less likely correct boundary, it will on average take longer to reach this boundary, which lowers the decision confidence since confidence decreases with elapsed time. Therefore, even though the decision confidence at both boundaries is the same for the given decision time, it will on average across decision times be lower for the a-priori non-preferred boundary, faithfully implementing this prior (see SI for a mathematical demonstration).

Our finding that a simple shift in the accumulation starting point is the Bayes-optimal strategy appears at odds with previous work that suggested that the optimal shift of the accumulator variable *z*(*t*) varies with time (18). This difference stems from a different implementation of the bias. While we have chosen an overall shift in the prior by its mean (Fig. 3c), an alternative implementation is to multiply *p*(*μ* ≥ 0) by *P*^+^, and *p*(*μ* < 0) by 1 − *P*^+^ (Fig. 3d), again resulting in *P*^+^ = *p*(*μ* ≥ 0). A consequence of this difference is that the associated shift of the posterior belief of *μ* ≥ 0 in the evidence accumulation space becomes time-dependent. Then, the optimal choice at a time-invariant boundary in that space might change over time (Fig. 3d). Furthermore, un-doing this shift to regain a unique association between boundaries and choices not only requires a shifted accumulation starting point, but additionally a time-dependent additive signal (*C*_0_(*P*^+^, *t*) in Fig. 3d; SI), as was proposed in (18). Which of the two approaches is more adequate depends on how well it matches the prior implicit in the task design. Our approach has the advantage of a simpler mechanistic implementation, as well as yielding a simple extension to the previously derived learning rule. How learning prior biases in the framework of (18) could be achieved remains unclear (but see (33)).

### Sequential choice dependencies due to continuous weight tracking

In every-day situations, no two decisions are made under the exact same circumstances. Nonetheless, we need to be able to learn from the outcome of past choices to improve future ones. A common assumption is that past choices become increasingly less informative about future choices over time. One way to express this formally is to assume that the world changes slowly over time – and that our aim is to track these changes. By ‘slow’ we mean that we can consider it constant over a single trial but that it is unstable over the course of an hour-long session. We implemented this tracking of the moving world, as in Fig. 2b, by slowly allowing the weights mapping evidence to decisions to change. With such continuously changing weights, weight learning never ends. Rather, the input weights are continuously adjusted to make correct choices more likely in the close future. After correct choices, this means that weights will be adjusted to repeat the same choice upon observing a similar input in the future. After incorrect choices, the aim is to adjust the weights to perform the opposite choice, instead. Our model predicts that, after an easy correct choice, in which confidence can be expected to be high, the weight adjustments are lower than after hard correct choices (see Fig. 1d top, green line). As a consequence, we would expect the model to be more likely to repeat the same choices after correct and hard, than after correct and easy trials.

To test this prediction, we relied on the same simulation to generate Fig. 2b to measured how likely the model repeated the same choice after correct decisions. Figure 4a illustrates that this repetition bias manifests itself in a shift of the psychometric curve that makes it more likely to repeat the previous choice. Furthermore, and as predicted, this shift is modulated by the difficulty of the previous choice and is stronger if the previous choice was easy (i.e., associated with a large |*μ*|; Fig. 4b). Therefore, if the decision maker expects to operate in a volatile, slowly changing world, our model predicts a repetition bias to repeat the same choices after correct decisions, and that this bias is stronger if the previous choice was easy.

**Figure 4.**
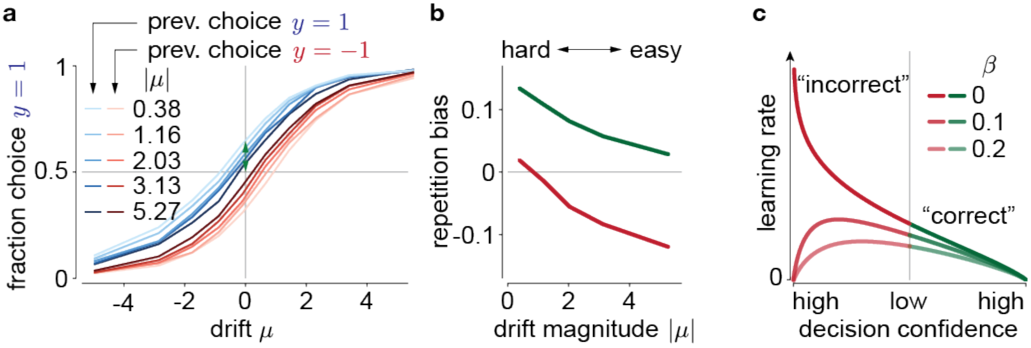
Sequential choice dependencies due to continuous learning, and effects of noisy feedback. Bayes-optimal learning in a slowly changing environment predicts sequential choice dependencies with the following pattern. (**a**) After hard, correct choices (low prev. |*μ*|; light colors), the psychometric curve is shifted towards repeating the same choice (blue/red = choice *y* = 1/−1). This shift decreases after easier, correct choices (high prev. |*μ*|; dark colors). (**b**) We summarize these tuning curve shifts in the repetition bias, which is the probability of repeating the same choice to a *μ* = 0 stimulus (example green arrow for *μ* = −0.38 in (**a**)). After correct/incorrect choices (green/red curve), this leads to a win-stay/lose-switch strategy. Only the win-stay strategy is shown in (**a**). (**c**) If choice feedback is noisy (inverted with probability *β*), the learning rate becomes overall lower. In particular for high-confidence choices with “incorrect” feedback, the learning rate becomes zero, as the learner trusts her choice more than the feedback.

### Unreliable feedback reduces learning

What would occur if choice feedback is less-than-perfectly reliable? For example, the feedback itself might not be completely trustworthy, or hard to interpret. We simulated this situation by assuming that the feedback is inverted with probability *β*. Here, *β* = 0 implies the so far assumed perfectly reliable feedback, and *β* = 1/2 makes the feedback completely uninformative. This change impacts how decision confidence modulates the learning rate (Fig. 4c) as follows. First, it reduces the overall magnitude of the correction, with weaker learning for higher feedback noise. Second, it results in no learning for highly confident choices that we are told are incorrect. In this case, one’s decision confidence overrules the unreliable feedback. This stands in stark contrast to the optimal learning rule for perfectly reliable feedback, in which case the strongest change to the current strategy ought to occur.

## Discussion

Diffusion models are applicable to model decisions that require some accumulation of evidence over time, which is almost always the case in natural decisions. We extended previous work on the normative foundations of these models to more realistic situations in which the sensory evidence is encoded by a population of neurons, as opposed to just two neurons, as has been typically assumed in previous studies. We have focused on normative and mechanistic models for learning the weights from the sensory neurons to the decision integrator without additionally adjusting the decision boundaries, as weight learning is a problem that needs to be solved even if the decision boundaries are optimized at the same time.

From the Bayesian perspective, weight learning corresponds to finding the weight posterior given the provided feedback, and resulted in an approximate learning rule whose learning rate was strongly modulated by decision confidence. It suppressed learning after high-confidence correct decisions, supported learning for uncertain decisions irrespective of their correctness, and promoted strong change of the combination weights after wrong decisions that were made with high confidence (Fig. 1d). Evidence for such confidence-based learning has already been identified in human experiments (34), but not in a task that required the temporal accumulation of evidence in individual trials. Indeed, as we have previously suggested (22), such a modulation by decision confidence should arise in all scenarios of Bayesian learning in N-AFC tasks in which the decision maker only receives feedback about the correctness of their choices, rather than being told which choice would have been correct. In the 2-AFC task we have considered, being told that one’s choice was incorrect automatically reveals that the other choice was correct, making the two cases coincide. Moving from one-dimensional to higher-dimensional inputs requires performing the accumulation of evidence for each input dimension separately (Fig. 1b; Eqs. (6) & (12) require ***x***(*t*) rather than only ***w***^a^***x***(*t*)), even if triggering choices only requires a linear combination of ***x***(*t*). This is because uncertain input weights require keeping track of how each input dimension contributed to the particle crossing the decision boundary in order to correctly improve these weights upon feedback (i.e., proper credit assignment). The multi-dimensional evidence accumulation predicted by our work arises naturally if inputs encode full distributions across the task-relevant variables, such as in linear probabilistic population codes (35) that trigger decisions by bounding the pooled activity of all units that represent the accumulated evidence (36).

Continual weight learning predicts sequential choice dependencies that make the repetition of a previous, correct choice more likely, in particular if this choice was difficult (Fig. 4). Thus, based on assuming a volatile environment that promotes a continual adjustment of the decision-making strategy, we provide a rational explanation for sequential choice dependencies that are frequently observed in both humans and animals (e.g., 37, 38). In rodents making decisions in response to olfactory cues we have furthermore confirmed that these sequential dependencies are modulated by choice difficulty, and that the exact pattern of this modulation depends on the stimulus statistics, as predicted by our theory (39) (but consistency with (40) unclear).

Lastly, we have clarified how prior biases ought to impact Bayes-optimal decision-making in diffusion models. Extending the work of Hanks et al. (18), we have demonstrated that the exact mechanisms to handle these biases depend on the specifics of how these biases are introduced through the task design. Specifically, we have suggested a variant that simplifies these mechanisms and the learning of this bias. This variant predicts that the evidence accumulation offset, that has previously been suggested to be time-dependent, to become independent of time, and it would be interesting to see if LIP activity of monkeys performing the random-dot motion task, as recorded by Hanks et al. (but see (41)), would change accordingly.

## Materials and Methods

We here provide an outline of the framework and its results. Detailed derivations are provided in the SI.

### Bayesian decision-making with one and multi-dimensional diffusion models

We assume the latent state to be drawn from 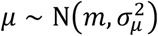, and the momentary evidence in each time step *δt* to provide information about this latent state by *δz*_*i*_|*μ* ∼ N(*μδt*, *δt*). The aim is to infer the sign of *μ*, and choose *y* = 1 if *μ* ≥ 0, and *y* = −1 otherwise. After having observed this evidence for some time *t* ≡ *nδt*, the posterior *μ* given all observed evidence *δz*_1:n_ is by Bayes’ rule given by

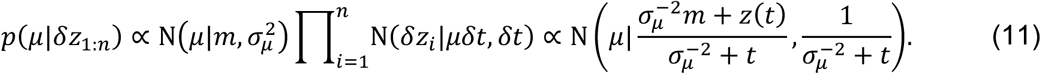

In the above, all proportionalities are with respect to *μ*, and we have defined 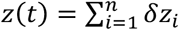 and have used 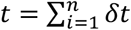. How to find the posterior belief for about *μ*’s sign with *m* = 0 is described around Eq. (1).

We extend diffusion models to multi-dimensional inputs with momentary evidence δ***x***_*i*_|*μ*, ***w*** ∼ *N*((***a**μ* + ***b***)*δt*, **Σ***δt*), with ***a***, ***b*** and **Σ** chosen such that ***w***^*T*^***x***(*t*)|*μ* = *z*(*t*)|*μ* ∼ *N*(*μt*, *t*), as before. The posterior over *μ* and *μ* ≥ 0 is the same as for the one-dimensional case, with *z*(*t*) replaced by ***w***^*T*^***x***(*t*). Definining 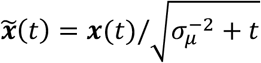, we find 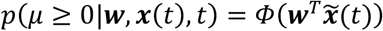. As *y* = 1 and *y* = −1 correspond to *μ* ≥ 0 and *μ* < 0, and *y* = 1 is only chosen if *p*(*μ* ≥ 0|***w***, ***x***(*t*), *t*) ≥ 1/2, the decision confidence for *m* = 0 at some boundary ***w***^*T*^***x***(*t*) = ±*θ*(*t*) is given by 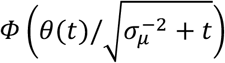. If input weights are unknown, and the decision maker holds belief ***w*** ∼ N(***μ***_*w*_, ***Σ***_*w*_) about these weights, the decision confidence needs to additionally account for weight uncertainty by marginalizing over ***w***, resulting in Eq. (5).

### Probabilistic and heuristic learning rules

We find the approximate posterior 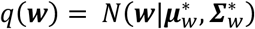 that approximates the target posterior *p* Eq. (4) by Assumed Density Filter (ADF). This requires minimizing the Kullback-Leiber divergence *KL*(*p*|*q*) (26, 29), resulting in Eq. (6) for the posterior mean, and

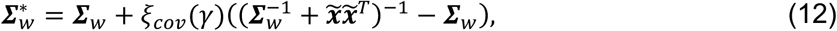

with learning rate modulators *ξ*_*w*_(*γ*) = N(*γ*|0,1)/Φ(*γ*) and *ξ*_*cov*_(*γ*) = *ξ*_*w*_(*γ*^2^ + *ξ*_*w*_(*γ*)*γ*, and where we have defined 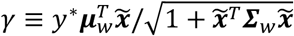, which is monotonic in the decision confidence, Eq. (5). Noisy choice feedback (Fig. 4c) changes the likelihood to assume reversed feedback with probability *β*, and follow the same procedure as above to derive the posterior moments (see SI). The ADF variant that only tracks the diagonal covariance elements assumes **Σ**_*w*_ to be diagonal, and only computes the diagonal elements of 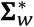. A second-order Taylor expansion of the log of Eq. (4) leads to update equations similar to Eqs. (6) and (12), but without the normalization by weight uncertainty (see SI for details). All heuristic learning rules are described in the main text.

We modeled non-stationary input weights by ***w***_*n*_|***w***_*n*−1_ ∼ N(***A******w***_*n*−1_ + ***b***, **Σ**_*d*_) after a decision in trial *n* − 1. This weight transition is taken into account by the probabilistic learning rules by setting the parameter priors to 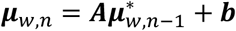 and 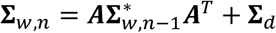. For stationary weights we have ***A*** = ***I***, ***b*** = 0, and **Σ**_*d*_ = **0**.

Bayes-optimal weight inference was for stationary weights performed by Gibbs sampling for Probit models, and for non-stationary weights by particle filtering (see SI).

### Simulation details

We used parameters ***a*** = ***w***/‖***w***‖^2^ and ***b*** = **0** for the momentary evidence *δ****x***. Its covariance **Σ** was generated to feature eigenvalues that drop exponentially from 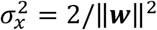 to zero until it reaches a constant 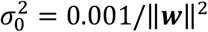 noise baseline, as qualitatively observed in neural populations. It additionally contains an eigenvector ***w*** with eigenvalue set to guarantee ***w***^***T***^**Σ*w*** = 1, limiting the information that *δ****x*** provides about *μ*. For non-stationary weights, all momentary evidence parameters are adjusted after each weight change (see SI). The diffusion model bounds ±*θ* were time-invariant and tuned to maximize the reward rate when using the correct weights. The reward rate is given by (*p*(*correct*) − *c*_*accum*_〈*t*〉)/(*t*_*iti*_ + 〈*t*〉), where averages where across trials, and we used evidence accumulation cost *c*_*accum*_ = 0.01 and inter-trial interval *t*_*iti*_ = 2*s*. We used 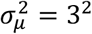 to draw *μ* in each trial, and drew ***w*** from ***w***~N(**1**, ***I***) before each trial sequence. For non-stationary weights, we re-sampled weight after each trial according to 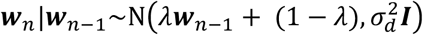, with decay factor *λ* = 1 − 0.01 and 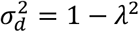 to achieve steady-state mean **1** and identity covariance.

To compare the weight learning performance of ADF to alternative models (Fig. 2a), we simulated 1,000 learning trials 5,000 times, and reported the reward rate per trial averaged across these 5,000 repetitions. To assess steady-state performance (Fig. 2b), we performed the same procedure with non-stationary weights, and reported reward rate averaged over the last 100 trials, and over 5,000 repetitions. The same 100 trials were used to compute the sequential choice dependencies in Fig. 4a/b. To simulate decision-making with diffusion models and uncertain weights, we used the current mean estimate 〈***w***〉 of the input weights to linearly combine the momentary evidence. The probabilistic learning rules were all independent of the specific choice of this estimate. The learning rate in Fig. 1d shows the pre-factor to 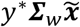 in Eq. (6) over decision confidence for a subsample of the last 10,000 trials of a single 15,000 trial simulation with non-stationary weights. For the Gibbs sampler, we drew 10 burn-in samples, followed by 200 samples in each trial. For the particle filter we simulated 1,000 particles.

## Acknowledgments

This work was supported by a James S. McDonnell Foundation Scholar Award (#220020462, JD), and grants from the NIMH (R01MH115554, JD), the Swiss National Science Foundation, www.snf.ch, (#31003A_143707 and #31003A_165831, AP), Champalimaud Foundation (ZFM), European Research Council (Advanced Investigator Grant 250334 & 67125, ZFM), Human Frontier Science Program (Grant RGP0027/2010, ZFM & AP), Simons Foundation (Grant 325057, ZFM & AP), and Fundação para a Ciência e a Tecnologia (AGM).

