## Supplemental Information for "Learning optimal decisions with confidence"

### Learning Optimal Decisions with Confidence
### Supplementary Information

Jan Drugowitsch, André G. Mendonça, Zachary F. Mainen and Alexandre Pouget

September 18, 2019

#### Supplementary Text

We here provide a self-consistent and extended derivation of all the results provided in the main text. We will use standard font for scalars, lower-case bold symbols for vectors, and upper-case bold symbols for matrices. All vectors are column vectors, unless transposed. $x \sim \mathcal{N}\left(\mu, \sigma^2\right)$ denotes that $x$ is a normal random variable with mean $\mu$ and variance $\sigma^2$.

#### Optimal decision making with high-dimensional momentary evidence

##### One-dimensional momentary evidence

Within each individual trial, we assume the latent state $\mu$ to be drawn from $\mu \sim \mathcal{N}\left(m, \sigma_\mu^2\right)$. $m = 0$ corresponds to the case of an unbiased prior for which $p(\mu \geq 0) = p(\mu < 0)$. In each small time step $n$ of size $\delta t$ from trial-onset at $t = 0$ (i.e., $n = 1$), the decision maker observes the momentary evidence $\delta z_n | \mu \sim \mathcal{N}\left(\mu \delta t, \delta t\right)$ that provides noisy information about the value of $\mu$. The decision-maker's aim is to infer the sign of $\mu$ from the sequence $\delta z_1, \delta z_2, \dots$ of momentary evidence, to make choice $y = 1$ (for $\mu \geq 0$) or $y = -1$ (for $\mu < 0$).

For Bayes-optimal choices, we find the posterior $p\left(\mu \geq 0 | \delta z_1, \delta z_2, \dots\right)$ in two steps. First, for $N$ pieces (i.e., $t = N\delta t$ seconds) of accumulated evidence, the posterior $\mu$ is give by Bayes' rule,

$$p\left(\mu | \delta z_{1:N}\right) \propto p\left(\mu\right) \prod_{n=1}^{N} p\left(\delta z_n | \mu\right) \propto e^{-\frac{\mu^2}{2}\left(\frac{1}{\sigma_\mu^2}\right) + \mu\left(\frac{m}{\sigma_\mu^2} + z\right)} \propto \mathcal{N}\left(\mu | \frac{\sigma_\mu^{-2} m + z}{\sigma_\mu^{-2} + t}, \frac{1}{\sigma_\mu^{-2} + t}\right), \tag{1}$$

where all proportionalities are with respect to $\mu$, and the second proportionality results from substituting the respective normal distributions, and defining $t = \sum_n \delta t$ and $z(t) = \sum_n \delta z_n$. This shows that the sufficient statistics of the posterior are $z(t)$ and $t$. For the second step we integrate this posterior over the non-negative half-line to find

$$p(\mu \geq 0 | z, t) = \int_0^\infty p(\mu | z, t) \mathrm{d}\mu = \Phi\left(\frac{\sigma_\mu^{-2} m + z}{\sqrt{\sigma_\mu^{-2} + t}}\right), \tag{2}$$

where $\Phi\left(\cdot\right)$ is the normal cumulative function. This posterior is more certain (i.e., closer to zero or one) for larger $|\sigma_\mu^{-2} m + z|$ and smaller times $t$.

Using the correspondence between $\mu \geq 0$ (and $\mu < 0$) and $y = 1$ (and $y = -1$), the fact that $1 - \Phi(a) = \Phi(-a)$, and $p\left(\mu < 0 | z, t\right) = 1 - p\left(\mu \geq 0 | z, t\right)$, the more generic posterior over $y$ is given by

$$p(y | z, t) = \Phi\left(y \frac{\sigma_\mu^{-2} m + z}{\sqrt{\sigma_\mu^{-2} + t}}\right). \tag{3}$$

This posterior captures both $y = 1$ and $y = -1$. If $y$ is the made decision, then the expression is the belief that this decision was correct, and hence the *decision confidence* [14].

So far we have assumed a prior over $\mu$ with arbitrary mean $m$. With this prior, the a-priori belief that $y = 1$ is correct is given by $P^+ \equiv p\left(\mu \geq 0\right) = \Phi\left(m / \sigma_\mu\right)$. The prior is thus unbiased for $m = 0$, in which case $P^+ = 1/2$. In this case, the posterior Eq. (3) prefers $y = 1$ for all $z > 0$ and $y = -1$ for all $z < 0$. Therefore, we can bound evidence accumulation from above and below by the (potentially time-dependent) $\pm\theta(t)$ to make Bayes-optimal choices. In particular, once $z$ reaches $\theta(t)$ (or $-\theta(t)$), it would trigger choice $y = 1$ (or $y = -1$). Observing that the unbounded accumulated evidence follows a Wiener process with drift $\mu$, that is, $z(t) | \mu \sim \mathcal{N}\left(\mu t, t\right)$, supports the use of drift-diffusion models for Bayes-optimal decision making. Biased priors, which we discuss in a later section, require additional attention to achieve Bayes-optimal choices.

#### High-dimensional momentary evidence

To move to $J$-dimensional momentary evidence $\delta \boldsymbol{x}$ while preserving parallels to the one-dimensional case, we assume that there exist some (for now, known) combination weights $\boldsymbol{w}$ such that $\delta z_n = \boldsymbol{w}^T \delta \boldsymbol{x}_n$. We achieve this by the generative model,

$$\delta \boldsymbol{x} | \mu \sim \mathcal{N}\left( \left( \boldsymbol{a}\mu + \boldsymbol{b} \right) \delta t, \boldsymbol{\Sigma} \delta t \right), \tag{4}$$

for vectors $\boldsymbol{a}$ and $\boldsymbol{b}$ that satisfy $\boldsymbol{a}^T \boldsymbol{w} = 1$ and $\boldsymbol{b}^T \boldsymbol{w} = 0$, and a covariance matrix $\boldsymbol{\Sigma}$ for which $\boldsymbol{w}^T \boldsymbol{\Sigma} \boldsymbol{w} = 1$. With these properties it becomes easy to show that $\boldsymbol{w}^T \delta \boldsymbol{x} | \mu \sim \mathcal{N}\left( \mu \delta t, \delta t \right)$, as required. We will discuss our specific choices for $\boldsymbol{a}$, $\boldsymbol{b}$, and $\boldsymbol{\Sigma}$ for the simulations shown in the main text further below.

Using the same steps as before, we find the posterior $\mu$ given $N$ steps (i.e., $t = N\delta t$ seconds) of momentary evidence to be given by

$$p\left( \mu | \delta \boldsymbol{x}_{1:N}, \boldsymbol{w} \right) = \mathcal{N}\left( \frac{\sigma_\mu^{-2} m + \boldsymbol{w}^T \boldsymbol{x}}{\sigma_\mu^{-2} + t}, \frac{1}{\sigma_\mu^{-2} + t} \right), \tag{5}$$

where we have defined the accumulated evidence $\boldsymbol{x}(t) = \sum_n \delta \boldsymbol{x}_n$. The posterior over $\mu \geq 0$ is correspondingly given by

$$p\left( \mu \geq 0 | \boldsymbol{x}, t, \boldsymbol{w} \right) = \Phi\left( \frac{\sigma_\mu^{-2} m + \boldsymbol{w}^T \boldsymbol{x}}{\sqrt{\sigma_\mu^{-2} + t}} \right). \tag{6}$$

Both expressions differ from the one-dimensional case by replacing $z$ by $\boldsymbol{w}^T \boldsymbol{x}$. Expressed as a posterior over $y$, the above turns into

$$p\left( y | \boldsymbol{x}, t, \boldsymbol{w} \right) = \Phi\left( y \frac{\sigma_\mu^{-2} m + \boldsymbol{w}^T \boldsymbol{x}}{\sqrt{\sigma_\mu^{-2} + t}} \right). \tag{7}$$

If $y$ is the made decision, the above is again the decision confidence. Note that the unbounded accumulated evidence follows the multi-dimensional drifting Wiener process, $\boldsymbol{x}(t) | \mu \sim \mathcal{N}\left( \left( \boldsymbol{a}\mu + \boldsymbol{b} \right) t, \boldsymbol{\Sigma} t \right)$, whose $\boldsymbol{w}$-weighted linear combination reduces to the same one-dimensional process $\boldsymbol{z}(t) = \boldsymbol{w}^T \boldsymbol{x}(t) \sim \mathcal{N}\left( \mu t, t \right)$ as before.

Assuming again an unbiased prior, $m = 0$, Bayes-optimal decisions are by the same logic as for the one-dimensional case cast by the boundaries $\pm\theta(t)$ on $\boldsymbol{w}^T \boldsymbol{x}$. Here, the positive (negative) boundary correspond to choice $y = 1$ ($y = -1$). We will discuss Bayes-optimal choices for biased priors in a later section.

#### If difficulty $|\mu|$ varies across trials, the decision confidence at a constant decision boundary drops over time

As the previous sections have shown, the decision confidence is the same for one- and high-dimensional momentary evidence as long as the decision boundary is on $z(t)$ and $\boldsymbol{w}^T \boldsymbol{x}(t)$, respectively. Furthermore, for time-invariant decision boundaries, $\pm\theta(t) = \pm\theta$, this decision confidence drops as a function of time. Here we show that this drop is a general property of symmetric priors over $\mu$ for which the difficulty $|\mu|$ can vary across trials, that extends beyond the Gaussian $p(\mu)$ we assume in other parts of this supplement. To show this, let us redefine $p(\mu)$ — in this section only — to be (as in [6]) given by

$$p(\mu) = \sum_{i=1}^{L} \frac{p_i}{2} \left( \delta(\mu - \mu_i) + \delta(\mu + \mu_i) \right), \tag{8}$$

which features $L$ point masses at $\pm\mu_1, \pm\mu_2, \dots, \pm\mu_L$, each weighted by $p_i/2$, and where we have assumed positive $p_i$ that satisfy $\sum_i p_i = 1$. Furthermore, without loss of generality, we assume the $\mu_i$'s to be positive, ordered, and unique, that is $0 < \mu_1 < \mu_2 < \dots < \mu_L$. Here, we disallow $\mu_1 = 0$ for notational convenience, but our argument can be easily extended to include this possibility. Assuming the same one-dimensional momentary evidence as before, $\delta z_n | \mu \sim \mathcal{N}\left( \mu \delta t, \delta t \right)$, it follows from Bayes' rule that

$$p\left( \mu = \mu_i | z, t \right) = \frac{p_i e^{-\frac{t}{2}\mu_i^2 + z\mu_i}}{\sum_j p_j e^{-\frac{t}{2}\mu_j^2} \left( e^{z\mu_j} + e^{-z\mu_j} \right)}. \tag{9}$$

Therefore, the belief that $\mu \geq 0$ ($y = 1$) at the upper boundary $z = \theta$ is given by

$$p\left( y = 1 | z = \theta, t \right) = \sum_i p\left( \mu = \mu_i | z = \theta, t \right) = \frac{\sum_i p_i e^{-\frac{t}{2}\mu_i^2 + \theta\mu_i}}{\sum_j p_j e^{-\frac{t}{2}\mu_j^2} \left( e^{\theta\mu_j} + e^{-\theta\mu_j} \right)}. \tag{10}$$

For our symmetric prior, this belief at the upper boundary equals the decision confidence at both boundaries. Therefore, we will use it as a proxy for decision confidence.

In what follows, we will show that this belief is a mixture of two components. The first is the belief that $\mu = \mu_i$ given some fixed difficulty, $\mu \in \{-\mu_i, \mu_i\}$, and the second is the probability that this is indeed the current difficulty. The first part turns out to be independent of time, whereas the second changes. In particular, as we will show, the probability that the difficulty is high (i.e., that $|\mu|$ is small) increases over time, resulting in a re-weighting of the per-difficulty beliefs. This re-weighting causes the overall belief to drop, as we argue in the main text.

Mathematically, this mixture can be written as

$$p\left(y=1|z=\theta,t\right) = \sum_i p\left(\mu=\mu_i|z=\theta,t\right) = \sum_i p\left(\mu=\mu_i|z=\theta,t,\mu=\pm\mu_i\right) p\left(\mu=\pm\mu_i|z=\theta,t\right). \tag{11}$$

In the right-most sum, the first probability is the per-difficulty belief $g_i$ for assumed difficulty $\mu_i$, and the second is the probability that $\mu_i$ is indeed the current difficulty. Both follow from (9), and are given by

$$g_i \equiv p\left(\mu=\mu_i|z=\theta,t,\mu=\pm\mu_i\right) = \frac{e^{\theta\mu_i}}{e^{\theta\mu_i}+e^{-\theta\mu_i}} = \frac{1}{1+e^{-2\theta\mu_i}}, \tag{12}$$

$$p\left(\mu=\pm\mu_i|z=\theta,t\right) = \frac{p_i e^{-\frac{t}{2}\mu_i^2}\left(e^{\theta\mu_i}+e^{-\theta\mu_i}\right)}{\sum_j p_j e^{-\frac{t}{2}\mu_j^2}\left(e^{\theta\mu_j}+e^{-\theta\mu_j}\right)} = \frac{w_i(t)}{\sum_j w_j(t)}, \tag{13}$$

where we have defined $w_i(t) = p_i e^{-\frac{t}{2}\mu_i^2}\left(e^{\theta\mu_i}+e^{-\theta\mu_i}\right)$ as the unnormalized, time-dependent, per-difficulty weights. This allows us to write the overall belief as the weighted mixture

$$p\left(y=1|z=\theta,t\right) = \sum_i \frac{w_i(t)}{\sum_j w_j(t)} g_i, \tag{14}$$

which is a weighted mixture of per-difficulty beliefs, $g_i$, in which only the mixture weights are time-dependent. Note that the per-difficulty beliefs are strictly increasing in $\mu_i$, such that are also ordered, that is, $g_1 < g_2 < \cdots < g_L$. Furthermore, increasing time by $\delta t > 0$ results in a drop in unnormalized weights, $w_i(t+\delta t) = a_i(\delta t) w_t(t)$ with $a_i(\delta t) = e^{-\frac{\delta t}{2}\mu_i^2} \in \{0, 1\}$. This drop is larger for larger $\mu_i$, that is $a_1(\delta t) > a_2(\delta t) > \cdots > a_L(\delta t)$. Therefore, increasing time results in putting proportionality more weight on per-difficulty beliefs associated with lower $\mu_i$'s with lower associated $g_i$, such that the overall belief, and equally the decision confidence, drops.

Once we stop varying the difficulty (i.e., $L = 1$), the belief reduces to the per-difficulty belief $g_1$, which does not drop over time. Therefore, the only condition required for the decision confidence to drop at a time-invariant boundary is for the difficulty $|\mu|$ to vary across trials.

### Learning the input combination weight $w$ from choice feedback

So far we have assumed $\boldsymbol{w}$ to be known. Here we derive learning rules for $\boldsymbol{w}$ based on feedback on the correctness of a choice. Specifically, we assume that the decision maker accumulated evidence $\boldsymbol{x}$ for some time $t$ and (potentially, but not necessarily) made decision $y$, after which feedback about the correct choice $y^*$ is provided. Before evidence accumulation we assume the decision maker to hold belief $p(\boldsymbol{w})$ about the input combination weights $\boldsymbol{w}$. Our aim is to find the posterior $p(\boldsymbol{w}|\boldsymbol{x}, t, y^*)$ given all the available evidence. We focus here on a feedback after a single choice. The same principles apply to choices sequences, by turning the posterior after a choice into the prior for the subsequent choice.

The desired posterior can be found by Bayes' rule

$$p\left(\boldsymbol{w}|\boldsymbol{x},t,y^*\right) \propto p\left(y^*|\boldsymbol{x},t,\boldsymbol{w}\right) p(\boldsymbol{w}), \tag{15}$$

where the likelihood $p\left(y^*|\boldsymbol{x},t,\boldsymbol{w}\right)$ is conditional on all observed quantities, $\boldsymbol{x}$ and $t$, and some hypothetical weights $\boldsymbol{w}$, and specifies the probability that $y^*$ is the correct choice given these weights. This likelihood turns out to correspond to the previously derived decision-making posterior, Eq. (7), which is a normal cumulative function with argument linear in $\boldsymbol{w}$. In general, problems with such a likelihood function structure are known as *Probit regression*. Such problems don't yield solutions for which the posterior has the same functional form as the prior — which is a desirable property to support efficient Bayesian input weight learning across longer sequences of choices, and to gain insight into the learning rule. Therefore, we derive below different approximations to such Bayes-optimal learning.

All of the below assumes an unbiased prior over $\mu$ by fixing $m$ to $m = 0$. We can extend the below rules to also learn the prior bias $m$ by extending the accumulated evidence vector $\boldsymbol{x}$ by one element fixed to $\sigma_\mu^{-2}$, and the weight vector $\boldsymbol{w}$ by one element containing $m$. Learning this extended weight vector then correspond to simultaneously learning the input weight and the prior bias.

#### The marginal decision confidence

Before deriving approximate weight learning rules, let us consider the consequences of uncertain weights on the decision confidence $p(y|\boldsymbol{x}, t)$ with these weights marginalized out. To do so, we assume our prior weight belief to be normal, $\boldsymbol{w} \sim \mathcal{N}\left(\boldsymbol{\mu}_w, \boldsymbol{\Sigma}_w\right)$ with mean

$\boldsymbol{\mu}_w$ and covariance $\boldsymbol{\Sigma}_w$. Then, we find this marginal decision confidence by first finding the marginal posterior over $\mu$, which is given by

$$p(\mu|\boldsymbol{x}, t) = \int p(\mu|\boldsymbol{x}, t, \boldsymbol{w})p(\boldsymbol{w})\mathrm{d}\boldsymbol{w} = \mathcal{N}\left(\frac{\boldsymbol{\mu}_w^T\boldsymbol{x}}{\sigma_\mu^{-2}+t}, \frac{1}{\sigma_\mu^{-2}+t} + \frac{\boldsymbol{x}^T\boldsymbol{\Sigma}_w\boldsymbol{x}}{\left(\sigma_\mu^{-2}+t\right)^2}\right), \tag{16}$$

were we have used Eq. (5) with $m = 0$. We find the marginal decision confidence $p(y|\boldsymbol{x}, t)$ by integrating the above over the non-negative halfline, which results after some simplification in

$$p(y|\boldsymbol{x}, t) = \Phi\left(y\frac{\boldsymbol{\mu}_w^T\boldsymbol{x}}{\sqrt{\sigma_\mu^{-2}+t+\boldsymbol{x}^T\boldsymbol{\Sigma}_w\boldsymbol{x}}}\right) = \Phi\left(y\frac{\boldsymbol{\mu}_w^T\tilde{\boldsymbol{x}}}{\sqrt{1+\tilde{\boldsymbol{x}}^T\boldsymbol{\Sigma}_w\tilde{\boldsymbol{x}}}}\right), \tag{17}$$

where we have defined $\tilde{\boldsymbol{x}} \equiv \boldsymbol{x}/\sqrt{\sigma_\mu^{-2}+t}$ for the second equality. Comparing this expression to Eq. (7) reveals the additional term $\boldsymbol{x}^T\boldsymbol{\Sigma}_w\boldsymbol{x}$ that lowers the overall posterior confidence (i.e., moving it towards 1/2) due to uncertainty in $\boldsymbol{w}$. If $y$ is the made choice, the above is the decision confidence that takes into account weight uncertainty.

#### Weight learning by Assumed Density Filtering

Assumed Density Filtering (ADF; [12, 10, 8, 3]) approximates the posterior by assuming a particular functional form of the approximate posterior $q(\boldsymbol{w}|\boldsymbol{x}, t, y^*)$ and finding the parameters of this functional form by minimizing the Kullback-Leiber divergence $\mathrm{KL}\left(p\left(\boldsymbol{w}|\boldsymbol{x}, t, y^*\right) \| q\left(\boldsymbol{w}|\boldsymbol{x}, t, y^*\right)\right)$ between the true posterior and its approximation. To minimize this divergence we again assume a normally distributed prior $\boldsymbol{w} \sim \mathcal{N}\left(\boldsymbol{\mu}_w, \boldsymbol{\Sigma}_w\right)$ with mean $\boldsymbol{\mu}_w$ and covariance $\boldsymbol{\Sigma}_w$. To support sequential choices, we assume the approximate posterior to also be normal, $q\left(\boldsymbol{w}|\boldsymbol{x}, t, y^*\right) = \mathcal{N}\left(\boldsymbol{w}|\boldsymbol{\mu}_w^*\boldsymbol{\Sigma}_w^*\right)$, with updated moments $\boldsymbol{\mu}_w^*$ and $\boldsymbol{\Sigma}_w^*$.

To find these updated moments, we use the fact that the KL-divergence is in our case minimized by matching the moments of the Gaussian sufficient statistics $\boldsymbol{w}$ and $\boldsymbol{w}\boldsymbol{w}^T$ [2]. For the source distribution, $p\left(\boldsymbol{w}|\boldsymbol{x}, t, y^*\right)$, these moments can be found by the gradients of the log-normalizing constant of this source distribution, $\nabla \log p(y^*|x, t)$ [2, 11], where we will use the already derived marginal likelihood $p(y|\boldsymbol{x}, t)$ in Eq. (17). Using these principles, the updated moments of the approximate posterior can be found by

$$\boldsymbol{\mu}_w^* = \boldsymbol{\mu}_w + \boldsymbol{\Sigma}_w\nabla_{\mu_w}\log p(y^*|\boldsymbol{x}, t), \tag{18}$$

$$\boldsymbol{\Sigma}_w^* = \boldsymbol{\Sigma}_w - \boldsymbol{\Sigma}_w\left(\nabla_{\mu_w}\log p(y^*|\boldsymbol{x}, t)\left(\nabla_{\mu_w}\log p(y^*|\boldsymbol{x}, t)\right)^T - 2\nabla_{\Sigma_w}\log p(y^*|\boldsymbol{x}, t)\right)\boldsymbol{\Sigma}_w. \tag{19}$$

The required gradients are given by

$$\nabla_{\mu_w}\log p(y^*|\boldsymbol{x}, t) = \xi_w(\gamma)y^*\frac{\tilde{\boldsymbol{x}}}{\sqrt{1+\tilde{\boldsymbol{x}}^T\boldsymbol{\Sigma}_w\tilde{\boldsymbol{x}}}}, \tag{20}$$

$$\nabla_{\Sigma_w}\log p(y^*|\boldsymbol{x}, t) = -\xi_w(\gamma)y^*\frac{\boldsymbol{\mu}_w^T\tilde{\boldsymbol{x}}}{2\sqrt{1+\tilde{\boldsymbol{x}}^T\boldsymbol{\Sigma}_w\tilde{\boldsymbol{x}}}^3}\tilde{\boldsymbol{x}}\tilde{\boldsymbol{x}}^T, \tag{21}$$

(22)

where $\xi_w(\gamma)$ is given by

$$\xi_w(\gamma) = \frac{\partial}{\partial\gamma}\log\Phi(\gamma) = \frac{\mathcal{N}\left(\gamma|0, 1\right)}{\Phi(\gamma)}, \tag{23}$$

and we have defined $\gamma$ as

$$\gamma \equiv y^*\frac{\boldsymbol{\mu}_w^T\tilde{\boldsymbol{x}}}{\sqrt{1+\tilde{\boldsymbol{x}}^T\boldsymbol{\Sigma}_w\tilde{\boldsymbol{x}}}}. \tag{24}$$

Overall, this leads to the moments update equations,

$$\boldsymbol{\mu}_w^* = \boldsymbol{\mu}_w + y^*\frac{\xi_w(\gamma)}{\sqrt{1+\tilde{\boldsymbol{x}}^T\boldsymbol{\Sigma}_w\tilde{\boldsymbol{x}}}}\boldsymbol{\Sigma}_w\tilde{\boldsymbol{x}}, \tag{25}$$

$$\boldsymbol{\Sigma}_w^* = \boldsymbol{\Sigma}_w + \xi_{cov}(\gamma)\left(\left(\boldsymbol{\Sigma}_w^{-1}+\tilde{\boldsymbol{x}}\tilde{\boldsymbol{x}}^T\right)^{-1} - \boldsymbol{\Sigma}_w\right), \tag{26}$$

where the covariance learning rate is given by

$$\xi_{cov}(\gamma) = \xi_w(\gamma)^2 + \xi_w(\gamma)\gamma. \tag{27}$$

As illustrated in Fig. 1, both mean and covariance updates are modulated by the marginal decision confidence in the feedback, $y^*$, given by Eq. (17). To see how $\xi_w$ and $\xi_{cov}$ are a function of the marginal decision confidence about the actual choice $y$ (rather than the feedback $y^*$) let us first focus on correct choices. For correct choices, $y = y^*$, such that the marginal decision confidence about $y$ equals that of $y^*$, that is, $p(y|\boldsymbol{x}, t) = p(y^*|\boldsymbol{x}, t)$. Furthermore, by the definition of $p(y^*|\boldsymbol{x}, t)$, it can be written as $p(y^*|\boldsymbol{x}, t) = \Phi(\gamma)$

(where $\gamma$ is defined in Eq. (24)). This function is strictly increasing in $\gamma$, such that small/large $\gamma$'s corresponds to low/high confidence. Therefore, as $\xi_w(\gamma)$ and $\xi_{cov}(\gamma)$ are functions of only $\gamma$, they are in turn functions of the marginal decision confidence $p(y|\boldsymbol{x}, t)$.

For incorrect choices we have $y \neq y^*$, such that $p(y|\boldsymbol{x}, t) = 1 - p(y^*|\boldsymbol{x}, t) = 1 - \Phi(\gamma)$, which is strictly decreasing in $\gamma$. Therefore, we can again assign a unique decision confidence $p(y|\boldsymbol{x}, t)$ to each $\gamma$, such that $\xi_w(\gamma)$ and $\xi_{cov}(\gamma)$ are again functions of the decision confidence about the made decision $y$.

#### Assumed Density Filtering with a diagonal covariance matrix

The above update equations require tracking of the full covariance matrix, making these updates scale badly with the size of the input space, $J$, and require non-local interactions. To find alternative, local update equations, we here assume that both the prior covariance, as well as the approximate posterior covariance are given by diagonal matrices, given by $\boldsymbol{\Sigma}_w = \mathrm{diag}\left(\sigma^2_{w,1}, \dots, \sigma^2_{w,k}\right)$ and $\boldsymbol{\Sigma}^*_w = \mathrm{diag}\left(\sigma^{2*}_{w,1}, \dots, \sigma^{2*}_{w,k}\right)$. Following the same derivation as before, this leads to the update equations

$$\mu^*_{w,i} = \mu_{w,i} + y^* \frac{\xi_w(\gamma)}{\sqrt{1 + \sum_j \sigma^2_{w,j} \tilde{x}^2_j}} \sigma^2_{w,i} \tilde{x}_i, \tag{28}$$

$$\sigma^{2*}_{w,i} = \sigma^2_{w,i} - \xi_{cov}(\gamma) \frac{\sigma^4_{w,i} \tilde{x}_i}{\sqrt{1 + \sum_j \sigma^2_{w,j} \tilde{x}^2_j}} \tag{29}$$

where $\mu_{w,i}$ and $\mu^*_{w,i}$ are the $i$th element of $\boldsymbol{\mu}_w$ and $\boldsymbol{\mu}^*_w$, respectively, and $\gamma$ is given by

$$\gamma = y^* \frac{\boldsymbol{\mu}^T_w \tilde{\boldsymbol{x}}}{\sqrt{1 + \sum_j \sigma^2_{w,j} \tilde{x}^2_j}}. \tag{30}$$

Thus, other than a global divisive normalization and the marginal decision confidence-related term $\gamma$, all updates are local.

#### Approximating the weight posterior by a second-order Taylor series

An simpler alternative to ADF that also yields a normally distributed approximate posterior is to approximate the true log-posterior, $\log p(\boldsymbol{w}|\boldsymbol{x}, t, y^*)$ by a second-order Taylor series in $\boldsymbol{w}$ around $\boldsymbol{w} = \boldsymbol{\mu}_w$. The relevant terms in this log-posterior are

$$\log p(\boldsymbol{w}|\boldsymbol{x}, t, y^*) = \Phi\left(y^* \boldsymbol{w}^T \tilde{\boldsymbol{x}}\right) - \frac{1}{2} \boldsymbol{w}^T \boldsymbol{\Sigma}^{-1}_w \boldsymbol{w} + \boldsymbol{w}^T \boldsymbol{\Sigma}^{-1}_w \boldsymbol{\mu}_w + \mathrm{const.} \tag{31}$$

The required gradient and Hessian are

$$\nabla_w \log p(\boldsymbol{w}|\boldsymbol{x}, t, y^*)\big|_{w=\mu_w} = \xi_w(\gamma) y^* \tilde{\boldsymbol{x}}, \tag{32}$$

$$\nabla\nabla_w \log p(\boldsymbol{w}|\boldsymbol{x}, t, y^*)\big|_{w=\mu_w} = -\boldsymbol{\Sigma}^{-1}_w - \xi_{cov}(\gamma) \tilde{\boldsymbol{x}} \tilde{\boldsymbol{x}}^T, \tag{33}$$

where $\xi_w(\cdot)$ and $\xi_{cov}(\cdot)$ are defined as for ADF, but $\gamma$ changes to $\gamma = y^* \boldsymbol{\mu}^T_w \tilde{\boldsymbol{x}}$. Using the above to find the second-order Taylor series and reading off the resulting posterior moments yields the moment updates

$$\boldsymbol{\mu}^*_w = \boldsymbol{\mu}_w + y^* \xi_w(\gamma) \boldsymbol{\Sigma}^*_w \tilde{\boldsymbol{x}}, \tag{34}$$

$$\boldsymbol{\Sigma}^*_w = \left(\boldsymbol{\Sigma}^{-1}_w + \xi_{cov}(\gamma) \tilde{\boldsymbol{x}} \tilde{\boldsymbol{x}}^T\right)^{-1}. \tag{35}$$

These have a similar form as for ADF, Eqs. (25) and (26), with the main difference that they are missing the normalization by $\sqrt{1 + \tilde{\boldsymbol{x}}^T \boldsymbol{\Sigma}_w \tilde{\boldsymbol{x}}}$. Given that this normalization modulates the moment update strength by the weight uncertainty, this implies that the update equations based on the second-order Taylor series will be less influenced by this uncertainty.

#### Assumed density filtering with noisy feedback

So far we have assumed the feedback $y^*$ to always be correct. We will now consider how ADF changes when the feedback itself is noisy. In particular, we assume that feedback is inverted with probability $\beta$, such that the weight likelihood given feedback $y^*$ becomes

$$p(y^*|\boldsymbol{x}, t, \boldsymbol{w}) = \beta \Phi\left(-y^* \boldsymbol{w}^T \tilde{\boldsymbol{x}}\right) + (1 - \beta) \Phi\left(y^* \boldsymbol{w}^T \tilde{\boldsymbol{x}}\right) = \beta + (1 - 2\beta) \Phi\left(y^* \boldsymbol{w}^T \tilde{\boldsymbol{x}}\right). \tag{36}$$

In this case, the marginal decision confidence about feedback $y^*$ becomes

$$p(y^*|\boldsymbol{x}, t, \beta) = \beta + (1 - 2\beta) \Phi\left(y^* \frac{\boldsymbol{\mu}^T_w \tilde{\boldsymbol{x}}}{\sqrt{1 + \tilde{\boldsymbol{x}}^T \boldsymbol{\Sigma}_w \tilde{\boldsymbol{x}}}}\right). \tag{37}$$

The gradients of the log marginal decision confidence thus become

$$\nabla_{\mu_w} \log p(y^*|\boldsymbol{x}, t, \beta) = \xi_{\beta,w}(\gamma) y^* \frac{\tilde{\boldsymbol{x}}}{\sqrt{1 + \tilde{\boldsymbol{x}}^T \boldsymbol{\Sigma}_w \tilde{\boldsymbol{x}}}}, \tag{38}$$

$$\nabla_{\Sigma_w} \log p(y^*|\boldsymbol{x}, t, \beta) = -\xi_{\beta,w}(\gamma) y^* \frac{\boldsymbol{\mu}_w^T \tilde{\boldsymbol{x}}}{2\left(1 + \tilde{\boldsymbol{x}}^T \boldsymbol{\Sigma}_w \tilde{\boldsymbol{x}}\right)^{3/2}} \tilde{\boldsymbol{x}} \tilde{\boldsymbol{x}}^T \tag{39}$$

with $\xi_{\beta,w}(\gamma)$ given by

$$\xi_{\beta,w}(\gamma) = \frac{\partial}{\partial \gamma} \log\left(\beta + (1-2\beta)\Phi(\gamma)\right) = \frac{(1-2\beta)\mathcal{N}\left(\gamma|0,1\right)}{\beta + (1-2\beta)\Phi(\gamma)} \tag{40}$$

and where $\gamma$ is, as for vanilla ADF, given by Eq. (24). Using again Eqs. (18) and (19) results in the update equations

$$\boldsymbol{\mu}_w^* = \boldsymbol{\mu}_w + y^* \frac{\xi_{\beta,w}(\gamma)}{\sqrt{1 + \tilde{\boldsymbol{x}}^T \boldsymbol{\Sigma}_w \tilde{\boldsymbol{x}}}} \boldsymbol{\Sigma}_w \tilde{\boldsymbol{x}}, \tag{41}$$

$$\boldsymbol{\Sigma}_w^* = \boldsymbol{\Sigma}_w + \xi_{\beta,cov}(\gamma) \left( \left( \boldsymbol{\Sigma}_w^{-1} + \tilde{\boldsymbol{x}} \tilde{\boldsymbol{x}}^T \right)^{-1} - \boldsymbol{\Sigma}_w \right), \tag{42}$$

with covariance learning rate

$$\xi_{\beta,cov}(\gamma) = \xi_{\beta,w}(\gamma)^2 + \xi_{\beta,w}(\gamma)\gamma. \tag{43}$$

This illustrates that the only impact of noisy feedback is on the update strength modulators, $\xi_{\beta,w}(\cdot)$ and $\xi_{\beta,cov}$. As shown in Fig. 1, these modulators become smaller for larger feedback noise. For high-confidence choices that the feedback flags as incorrect, $\xi_{\beta,cov}$ even becomes negative, indicating that uncertainty in $\boldsymbol{w}$ increases. This increase arises from approximate inference, as additional information in strictly Bayes-optimal inference cannot increase uncertainty, even if this information is (knowingly) noisy.

##### Alternative learning heuristics

Let us now discuss alternative heuristics that do not track a belief over $\boldsymbol{w}$, but instead update a point estimate. The first alternative is the delta rule that performs stochastic gradient descent on the sum of squared distances between the chosen decision boundary and the correct decision boundary. For the current choice, this squared distance is $\left(\boldsymbol{w}^T \boldsymbol{x}(t) - y^* \theta(t)\right)^2$, where $\boldsymbol{w}^T \boldsymbol{x}(t) \in \{-\theta(t), \theta(t)\}$ at decision time $t$ equals the chosen bound, and $y^* \theta(t) \in \{-\theta(t), \theta(t)\}$ is the boundary that would have led to the correct choice. Thus, the delta rule update is given by

$$\boldsymbol{w}^* = \boldsymbol{w} + \frac{\alpha}{2\theta(0)} \left( y^* \theta(t) - \boldsymbol{w}^T \boldsymbol{x} \right) \boldsymbol{x}, \tag{44}$$

where we have chosen to normalize the learning $\alpha$ by $2\theta(0)$ to make the update magnitude less dependent of the bound height. The residual in the above is either zero or $\pm 2\theta(t)$, such that the learning rule only makes adjustments to the weight estimate in case of incorrect choices.

The delta rule aims to minimize the probability that incorrect choices are made. In diffusion models this can be achieved by accumulating more evidence before reaching the decision boundary. This, in turn, can be accomplished by reducing the overall magnitude of $\boldsymbol{w}$. In particular for small learning rates, this is exactly what the delta rule does, leading to progressively smaller $\|\boldsymbol{w}\|$, and weight learning that does not converge in expectation. To work around this degeneracy, we introduced the normalized delta rule. This rule performs the update exactly like the standard delta rule, but subsequently adjusts the weight magnitude to match that of the true weights. It therefore needs access to the true weight's magnitude in each trial, making it a rule that has access to an oracle that other rules don't. Thus, it uses strictly more information than other rules, which needs to be kept in mind when comparing its performance to that of other rules.

As a last heuristic we considered performing stochastic gradient descent on the log-likelihood of the feedback, $\log p\left(y^*|\boldsymbol{x}, \boldsymbol{w}, t\right) = \log \Phi\left(y^* \boldsymbol{w}^T \tilde{\boldsymbol{x}}\right)$. Taking the gradient of this log-likelihood results in the learning rule

$$\boldsymbol{w}^* = \boldsymbol{w} - \alpha y^* \xi_w \left( y^* \boldsymbol{w}^T \tilde{\boldsymbol{x}} \right) \tilde{\boldsymbol{x}}, \tag{45}$$

where $\xi_w(\cdot)$ is defined as for ADF. Due to the inclusion of $\xi_w(\cdot)$, this rule modulates the update strength by decision confidence, unlike the normalized delta rule above. It differs from probabilistic learning rules in that it uses a fixed learning rate $\alpha$, instead of a learning rate modulation by a current estimate of the certainty about $\boldsymbol{w}$.

#### Tracking non-stationary combination weights

So far we have assumed the true weights, underlying the generation of the momentary evidences, $\delta\boldsymbol{x}$, to be stationary, allowing us to use a sequence $\boldsymbol{x}$'s, $t$'s, and $y^*$'s to learn successively better posteriors over $\boldsymbol{w}$. In the ideal case (i.e., if we wouldn't use approximate inference), this would — after enough observations — lead to a very good approximation of the true $\boldsymbol{w}$. We now change this setup to assume that the true weights change slightly across successive trials, and the learner's task is to track these changes as well as possible. This implies that, as the weights are now a moving target, they can never be learned perfectly.

We model the non-stationary of the weights by a first-order autoregressive process. That is, we assume that the true weights $\boldsymbol{w}_n$ in trial $n$ depend on the true weights $\boldsymbol{w}_{n-1}$ in trial $n$ by

$$\boldsymbol{w}_n | \boldsymbol{w}_{n-1} \sim \mathcal{N}\left(\boldsymbol{A}\boldsymbol{w}_{n-1} + \boldsymbol{b}, \boldsymbol{\Sigma}_d\right), \tag{46}$$

where $\boldsymbol{A}$, $\boldsymbol{b}$ and $\boldsymbol{\Sigma}_d$ are parameters of the process.

Let us now consider a probabilistic learner that maintains belief $\boldsymbol{w}_n \sim \mathcal{N}\left(\boldsymbol{\mu}_{w,n}, \boldsymbol{\Sigma}_{w,n}\right)$ before observing $\boldsymbol{x}$, $t$, and $y^*$ in the $n$th trial. Despite the successive weight change across trials, the learner would first follow its standard learning rule (discussed above different approximations) to compute posterior parameters $\boldsymbol{\mu}^*_{w,n}$ and $\boldsymbol{\Sigma}^*_{w,n}$. This is followed by taking account of the weight change by updating its parameters according to

$$\boldsymbol{\mu}_{w,n+1} = \boldsymbol{A}\boldsymbol{\mu}^*_{w,n} + \boldsymbol{b}, \qquad \boldsymbol{\Sigma}_{w,n+1} = \boldsymbol{A}\boldsymbol{\Sigma}^*_{w,n}\boldsymbol{A}^T + \boldsymbol{\Sigma}_d. \tag{47}$$

These weights then act as a starting point, i.e., prior, for learning in the next trial. No other changes to the learning rules are required to take the non-stationarity of the combination weights into account.

### Sampling the Bayes-optimal posterior

Finding a tractable closed-form expression for the Bayes-optimal posterior over $\boldsymbol{w}$ is unfortunately impossible. However, we can approximate this posterior to almost arbitrary precision by drawing samples from this posterior. We will first discuss such sampling for stationary combination weights, in which case we can use Gibbs sampling.

#### Gibbs sampling for stationary weights

For Gibbs sampling, we assume prior $\boldsymbol{w} \sim \mathcal{N}\left(\boldsymbol{\mu}_0, \boldsymbol{\Sigma}_0\right)$, and observations $\boldsymbol{x}_n$, $t_n$, and $y^*_n$ in the $n$th trial. The aim is to, after $N$ trials, draw samples from $p\left(\boldsymbol{w}|\boldsymbol{x}_{1:N}, t_{1:N}, y^*_{1:N}\right)$. With the per-trial likelihood $p\left(y^*_n|\boldsymbol{x}_n, t_n, \boldsymbol{w}\right) = \Phi\left(y^*_n \boldsymbol{w}^T \tilde{\boldsymbol{x}}_n\right)$, this posterior is given by

$$p\left(\boldsymbol{w}|\boldsymbol{x}_{1:N}, t_{1:N}, y^*_{1:N}\right) \propto \mathcal{N}\left(\boldsymbol{w}|\boldsymbol{\mu}_0, \boldsymbol{\Sigma}_0\right) \prod_{n=1}^{N} \Phi\left(y^*_n \boldsymbol{w}^T \tilde{\boldsymbol{x}}_n\right). \tag{48}$$

The covariance of this posterior is given by

$$\boldsymbol{\Sigma}_w = \left(\boldsymbol{\Sigma}_0^{-1} + \sum_{n=1}^{N} \tilde{\boldsymbol{x}}_n \tilde{\boldsymbol{x}}_n^T\right)^{-1} \tag{49}$$

which can be efficiently updated with each successive trial by the Sherman-Morrison update. To sample from the posterior $\boldsymbol{w}$, we introduce the auxiliary variables $a_n \sim \mathcal{N}\left(y^*_n \boldsymbol{w}^T \tilde{\boldsymbol{x}}_n, 1\right)$ for each $n$, such that $a_n \geq 0$ for a choice consistent with $y^*_n$. Thus, for a fixed $\boldsymbol{w}$, we can draw $a_n$ according to

$$a_n \sim \mathcal{N}_{\geq 0}\left(y^*_n \boldsymbol{w}^T \tilde{\boldsymbol{x}}_n, 1\right), \tag{50}$$

where $\mathcal{N}_{\geq 0}$ denotes a draw from a truncated normal distribution, guaranteeing $a_n \geq 0$. With these samples, the posterior $\boldsymbol{w}$ is given by

$$\boldsymbol{w}|\boldsymbol{x}_{1:N}, t_{1:N}, y^*_{1:N}, a_{1:N} \sim \mathcal{N}\left(\boldsymbol{\Sigma}_w \left(\Sigma_0^{-1}\boldsymbol{\mu}_0 + \sum_{n=1}^{N} y^*_n \tilde{\boldsymbol{x}}_n a_n\right), \boldsymbol{\Sigma}_w\right). \tag{51}$$

Overall, Gibbs sampling consists in alternating between sampling $a_{1:N}$ and $\boldsymbol{w}$ until a sufficient number of $\boldsymbol{w}$-samples are drawn.

#### Particle filtering for non-stationary weights

Once the weights become non-stationary, particle filtering turns out to be a more efficient approach to posterior sampling. The aim is to approximate the sequential weight update

$$p\left(\boldsymbol{w}_n|\boldsymbol{x}_{1:n}, \boldsymbol{t}_{1:n}, y^*_{1:n}\right) \propto p\left(y^*_n|\boldsymbol{x}_n, t_n, \boldsymbol{w}_n\right) \int p\left(\boldsymbol{w}_n|\boldsymbol{w}_{n-1}\right) p\left(\boldsymbol{w}_{n-1}|\boldsymbol{x}_{1:n-1}, t_{1:n-1}, y^*_{1:n-1}\right) \mathrm{d}\boldsymbol{w}_{n-1}, \tag{52}$$

by using the particle approximation

$$p\left(\boldsymbol{w}_n|\boldsymbol{x}_{1:n}, \boldsymbol{t}_{1:n}, y^*_{1:n}\right) \approx \frac{1}{K} \sum_{k=1}^{K} \delta_{w_n^{(k)}}, \tag{53}$$

consisting of the $K$ particles $\left\{\boldsymbol{w}_n^{(1)}, \ldots, \boldsymbol{w}_n^{(K)}\right\}$. With this approximation, the above sequential update becomes

$$p\left(\boldsymbol{w}_n | \boldsymbol{x}_{1:n}, t_{1:n}, t_{1:n}^*\right) \overset{\propto}{\approx} \sum_k p\left(y_n^* | \boldsymbol{x}_n, t_n, \boldsymbol{w}_n\right) p\left(\boldsymbol{w}_n | \boldsymbol{w}_{n-1}^{(k)}\right). \tag{54}$$

We can sample from this posterior by an importance sampling re-sampling scheme in three steps. First, we draw $K$ samples $\tilde{\boldsymbol{w}}_n^{(k)}$ from a Gaussian proposal density

$$\tilde{\boldsymbol{w}}_n^{(k)} \sim \mathcal{N}\left(\boldsymbol{\mu}_w\left(\boldsymbol{w}_{n-1}^{(k)}\right), \boldsymbol{\Sigma}_w\left(\boldsymbol{w}_{n-1}^{(k)}\right)\right). \tag{55}$$

Second, we compute the importance sampling weights,

$$\lambda_n^{(k)} = \frac{p\left(y_n^* | \boldsymbol{x}_n, t_n, \tilde{\boldsymbol{w}}_n^{(k)}\right) p\left(\tilde{\boldsymbol{w}}_n^{(k)} | \boldsymbol{w}_{n-1}^{(k)}\right)}{\mathcal{N}\left(\boldsymbol{\mu}_w\left(\boldsymbol{w}_{n-1}^{(k)}\right), \boldsymbol{\Sigma}_w\left(\boldsymbol{w}_{n-1}^{(k)}\right)\right)}. \tag{56}$$

Third, we re-sample the $\boldsymbol{w}_n^{(k)}$'s from the $\tilde{\boldsymbol{w}}_n^{(k)}$'s with probabilities proportional to their respective weights, $\lambda_n^{(k)}$. To ensure efficiency of the procedure, the proposal density for each weight should be close to $p\left(y_n^* | \boldsymbol{x}_n, t_n, \boldsymbol{w}_n\right) p\left(\boldsymbol{w}_n | \boldsymbol{w}_{n-1}\right)$, appropriately normalized, which we achieve by computing the proposal moments $\boldsymbol{\mu}_w\left(\boldsymbol{w}_{n-1}^{(k)}\right)$ and $\boldsymbol{\Sigma}_w\left(\boldsymbol{w}_{n-1}^{(k)}\right)$ according to the ADF variant that assumes non-stationary combination weights.

### Relating learning through inference to learning through optimization

In all of the above we have treated learning as an inference problem, where we want to find the posterior weights given all of the observed evidence. Here, we address the parallels between inference and optimization in two ways. First, we will describe more general decision theoretical principles that highlight these parallels. Second, we will show explicitly how our learning problem can be formulated as an optimization problem.

#### Decision theoretic perspective

In decision theory, the Bayes-optimal decision rule is the rule that minimizes some expected loss [1]. In our case, we have defined the loss as the negative reward rate, which is the negative average number of correct decisions per unit time, across a long sequence of such decisions. Furthermore, we have tuned the diffusion model boundaries such that there exists an optimal set of weights $\boldsymbol{w}^*$ that maximize the reward rate, and thus minimize the loss. Formally, the loss function $L\left(\boldsymbol{w}^*, \boldsymbol{w}\right)$ returns this loss for a given action $\boldsymbol{w}$ (in our case a particular set of chosen weights) given some unobserved state of nature, $\boldsymbol{w}^*$ (in our case the set of weights that maximize the reward rate).

Given observations $X$ (here, all information gathered from past trials), the Bayes-optimal action is the one minimizing the posterior loss, that is

$$\underset{\boldsymbol{w}}{\operatorname{argmin}} \left\langle L(\tilde{\boldsymbol{w}}, \boldsymbol{w})\right\rangle_{p(\tilde{\boldsymbol{w}}|X)}, \tag{57}$$

where $p\left(\tilde{\boldsymbol{w}}|X\right)$ are the posterior weights given all past information. If we assume the loss to be approximately quadratic around $\tilde{\boldsymbol{w}}$, then it is (approximately) minimized by $\langle\tilde{\boldsymbol{w}}|\boldsymbol{X}\rangle$ [1]. This justifies computing the posterior to perform learning through inference, and the use of the posterior mean for decision-making, as used in the main text.

Using Bayes-optimal decision rules for decision making has several appealing properties. One of particular interest in relation to learning through optimization is that it is an admissible rule [1, Ch. 4, Th. 9]. Here, admissability is a concept from the frequentist school of decision theory, and specifies a (not necessarily unique) decision rule $\delta(\cdot)$ whose associated risk function $R(\boldsymbol{w}^*, \delta)$ is smallest among all possible decision rules and all possible states of nature $\boldsymbol{w}^*$. Here, the risk function is the expected loss for a given $\boldsymbol{w}^*$, with the expectation taken over possible observations $X$ given $\boldsymbol{w}^*$, that is $R(\boldsymbol{w}^*, \delta) = \left\langle L(\boldsymbol{w}^*, \delta(X))\right\rangle_{p(X|\boldsymbol{w}^*)}$. Therefore, the Bayes-optimal decision rule doesn't only minimize the expected loss under the posterior, but also the expected loss across different (frequentist) repetitions of the same "experiment", that is, different observations for the same state of nature $\boldsymbol{w}^*$, and does so across all possible states of nature. As a consequence, finding the posterior $\boldsymbol{w}$ through inference allows us to make decisions that (approximately) minimize the loss in multiple senses, which, in our case, maximizes the reward rate.

#### Explicit learning through optimization

Here we demonstrate for the stationary-weight case that our inference problem can be formulated as an optimization problem that aims at maximizing performance — here for simplicity measured as the probability of making correct choices. To do so, assume that, in each trial, the decision maker observes some $J$-dimensional momentary evidence $\delta x$ that relates to the underlying latent state $\mu$ by Eq. (4), as before. They accumulate this evidence into $\boldsymbol{x}(t)$, and at some point (e.g., when a decision boundary is reached) decide according to $y = \text{sign}\left(\boldsymbol{w}^T \boldsymbol{x}(t)\right)$, using some decision strategy weight parameters $\boldsymbol{w}$. Their aim is to optimize these weight parameters to maximize their probability of making correct choices.

To find the solution to this maximization problem, let us establish which weight parameters maximize the probability of making correct choices. For this, note that by Eq. (4), the accumulated evidence is distributed as

$$\boldsymbol{x}(t)|\mu^* \sim \mathcal{N}\left(\left(\boldsymbol{a}\mu^* + \boldsymbol{b}\right)t, \boldsymbol{\Sigma}t\right), \tag{58}$$

where $\mu^*$ is the (unobserved) latent state that determines the correct choice by $y^* = \text{sign}(\mu^*)$. As a consequence, $\boldsymbol{w}^T\boldsymbol{x}(t)/t$ is distributed as

$$\frac{\boldsymbol{w}^T\boldsymbol{x}(t)}{t}|\mu^* \sim \mathcal{N}\left(\boldsymbol{w}^T\boldsymbol{a}\mu^* + \boldsymbol{w}^T\boldsymbol{b}, \frac{1}{t}\boldsymbol{w}^T\boldsymbol{\Sigma}\boldsymbol{w}\right). \tag{59}$$

Recall that $\boldsymbol{a}$, $\boldsymbol{b}$ and $\boldsymbol{\Sigma}$ in Eq. (4) have been defined to satisfy $\boldsymbol{w}^{*T}\boldsymbol{a} = 1$, $\boldsymbol{w}^{*T}\boldsymbol{b} = 0$, and $\boldsymbol{w}^{*T}\boldsymbol{\Sigma}\boldsymbol{w}^* = 1$ for some particular $\boldsymbol{w}^*$. For these parameters, we thus have $\boldsymbol{w}^{*T}\boldsymbol{x}(t)/t \sim \mathcal{N}\left(\mu^*, t^{-1}\right)$, which provides the best estimate of $\mu^*$ (in the mean squared error sense [1]), that can in turn be used as a basis for decision-making.

To find $\boldsymbol{w}^*$ from an observed sequence of $(\boldsymbol{x}_1(t_1), t_1, y_1^*), (\boldsymbol{x}_2(t_2), t_2, y_2^*), \dots (\boldsymbol{x}_N(t_N), t_N, y_N^*)$, we can use maximum likelihood, which is consistent and asymptotically efficient. For the diffusion model, the likelihood of $\boldsymbol{w}$ for a particular choice $y$ is by Eq. (7) (using $m = 0$) given by $p\left(y|\boldsymbol{x}, t, \boldsymbol{w}\right) = \Phi\left(y\boldsymbol{w}^T\tilde{\boldsymbol{x}}\right)$, where $\tilde{\boldsymbol{x}} \equiv \boldsymbol{x}/\sqrt{t + \sigma_\mu^{-2}}$, as previously defined. Therefore, the maximum (log-)likelihood estimate for the observed sequence is given by

$$\hat{\boldsymbol{w}}_{ML} = \underset{\boldsymbol{w}}{\text{argmax}} \sum_{n=1}^{N} \log\Phi\left(y_n^*\boldsymbol{w}^T\tilde{\boldsymbol{x}}_n\right). \tag{60}$$

Finding this estimate is an optimization problem. For a small number of observations $N$, this optimization problem might be underdetermined. To avoid instabilities, we can additionally add a regularization term that penalizes too large $\|\boldsymbol{w}\|^2$, leading to

$$\hat{\boldsymbol{w}}_{ML,reg} = \underset{\boldsymbol{w}}{\text{argmax}} \left(-\lambda\boldsymbol{w}^T\boldsymbol{w} + \sum_{n=1}^{N} \log\Phi\left(y_n^*\boldsymbol{w}^T\tilde{\boldsymbol{x}}_n\right)\right), \tag{61}$$

where $\lambda > 0$ is some regularization parameter. Overall, this demonstrates how to formulate our learning problem as a an optimization problem. We have used this approach to formulate one of our heuristics, resulting in Eq. (45).

To see how this approach relates to learning through inference, compare the expression for $\hat{\boldsymbol{w}}_{ML,reg}$ to Eq. (48). As can be seen, with prior parameters $\boldsymbol{\mu}_0 = \boldsymbol{0}$ and $\boldsymbol{\Sigma}_0 = \lambda^{-1}\boldsymbol{I}$, $\hat{\boldsymbol{w}}_{ML,reg}$ finds the maximum of the Bayesian parameter posterior over $\boldsymbol{w}$, given by Eq. (48). In other words, it equals the maximum a-posteriori estimate. However, the optimization approach does not directly provide an estimate of the uncertainty in $\hat{\boldsymbol{w}}_{ML,reg}$. This makes it hard to form consistent sequential updates, in which uncertain weights should be updated more strongly than certain weights. More generally, formulating the learning problem as an optimization problem reduces our ability to interpret the resulting expressions. For example, we might not have been able to identify that the learning rate is modulated by decision confidence without the inference formulation. All of these points made us follow the learning-as-inference route instead.

### Implementing prior biases

So far we have assumed $P^+ \equiv p(\mu \geq 0) = 1/2$, making both $\mu \geq 0$ and $\mu < 0$ equally likely. Let us now consider how to consistently implement prior biases for which $P^+ \neq 1/2$. To do so, we will restrict our discussion to the one-dimensional momentary evidence $\delta z$. The high-dimensional momentary evidence case follows the same principles, and yields the same conclusions, but it notationally more burdensome.

With a *consistent* implementation of a prior bias we mean that we want to be able to choose a pair of arbitrary, potentially time-changing boundaries[1] $\pm\theta(t)$, each of which triggers a different Bayes-optimal choice. This requirement turns out to become critical.

Let us discuss two ways to implement biased priors in turn. The first corresponds to a shift in the mean of $p(\mu)$, while the second modulates the mass of $\mu \geq 0$ while keeping the shape of $p(\mu)$ otherwise unchanged (as in [9]).

#### Shifting the prior mean

If we assume prior $\mu \sim \mathcal{N}(m, \sigma_\mu)$, then $P^+ = \Phi(m/\sigma_\mu)$, such that $P^+ \neq 1/2$ if and only if $m \neq 0$. This is the case we have discussed further above (see Sec. "One-dimensional momentary evidence"), where we have found the posterior

$$p(\mu > 0|z, t) = \Phi\left(\frac{\sigma_\mu^{-2}m + z}{\sqrt{\sigma_\mu^{-2} + t}}\right). \tag{62}$$

[1] They might even follow different time-courses, without changing any of the discussed concepts. To keep notation simple, we won't consider this case.

Thus, the posterior is $p(\mu \geq 0|z,t) \geq 1/2$ if and only if $\sigma_\mu^{-2}m + z \geq 0$. This implies that Bayes-optimal decisions are determined by the sign of $\sigma_\mu^{-2}m + z$. As a result, we cannot simply bound the accumulated evidence $z$, as this might not guarantee a unique association between boundaries and Bayes-optimal choices. For example, consider some negative $m < 0$ and a positive $z < \sigma_\mu^{-2}|m|$ that has just reached the upper boundary $z = \theta(t)$. At this point we would intuitively make choice $y = 1$, corresponding to $\mu \geq 0$. However, as $\sigma_\mu^{-2}m + z < 0$, our expression for the posterior shows that $p(\mu \geq 0|z,t) < 1/2$, such that $y = -1$ would be the Bayes-optimal choice. This shows that bounding $z$ directly can in some cases violate the boundary - choice correspondence.

We can regain this correspondence by instead bounding $\tilde{z}(t) = \sigma_\mu^{-2}m + z(t)$, which, by definition, starts at $\tilde{z}(0) = \sigma_\mu^{-2}m$. For this new accumulation variable it is easy to see that $p(\mu \geq 0|\tilde{z},t) \geq 1/2$ if and only if $\tilde{z} \geq 0$, thus restoring the boundary - choice correspondence.

#### Directly modulating $p(\mu \geq 0)$

An alternative approach to introducing a biased prior, which was taken in [9], is to boost one half of $p(\mu)$, while modulating down the other half,

$$p(\mu) = 2\mathcal{N}\left(\mu|0,\sigma_\mu^2\right)\begin{cases} P^+ & \text{if } \mu \geq 0, \\ 1-P^+ & \text{otherwise,}\end{cases} \tag{63}$$

ensuring again $p(\mu \geq 0) = P^+$. This prior, and the corresponding solution, has previously been investigated by [9].

This choice of prior results in the posterior over $\mu$,

$$p(\mu|z,t) \propto \mathcal{N}\left(\mu|\frac{z}{\sigma_\mu^{-2}+t}, \frac{1}{\sigma_\mu^{-2}+t}\right)\begin{cases} P^+ & \text{if } \mu \geq 0, \\ 1-P^+ & \text{otherwise .}\end{cases} \tag{64}$$

Adding the normalization constant and integrating the above over all $\mu \geq 0$ results in the posterior

$$p\left(\mu \geq 0|z,t\right) = \frac{P^+\Phi\left(\frac{z}{\sqrt{\sigma_\mu^{-2}+t}}\right)}{P^+\Phi\left(\frac{z}{\sqrt{\sigma_\mu^{-2}+t}}\right) + (1-P^+)\left(1-\Phi\left(\frac{z}{\sqrt{\sigma_\mu^{-2}+t}}\right)\right)}. \tag{65}$$

This posterior is $p\left(\mu \geq 0|z,t\right) \geq 1/2$, and thus promotes choice $y = 1$, if

$$\log \frac{\Phi\left(\frac{z}{\sqrt{\sigma_\mu^{-2}+t}}\right)}{1-\Phi\left(\frac{z}{\sqrt{\sigma_\mu^{-2}+t}}\right)} \geq \log\frac{1-P^+}{P^+}, \tag{66}$$

that is, if the log-odds provided by the accumulated evidence exceeds that of the prior log-odds for $\mu < 0$. For the same accumulator value $z$, the evidence log-odds drops to zero over time. As a result, it might be that the Bayes-optimal choice at the same boundary changes over time, thus violating the boundary - decision correspondence.

When compared to the previous section, the way the prior impacts the posterior is more complex. This makes recovering the boundary - decision correspondence more complex. The aim is to find a $C_2(P^+,t)$ such that $\tilde{z}(t) = z(t) + C_2(P^+,t)$ determines Bayes-optimal decisions by its sign alone. This can be achieved by

$$C_2(P^+,t) = \sqrt{\sigma_\mu^{-2}+t}\,\Phi^{-1}\left(\frac{P^+\Phi\left(\frac{z}{\sqrt{\sigma_\mu^{-2}+t}}\right)}{P^+\Phi\left(\frac{z}{\sqrt{\sigma_\mu^{-2}+t}}\right) + (1-P^+)\Phi\left(\frac{-z}{\sqrt{\sigma_\mu^{-2}+t}}\right)}\right) - z(t), \tag{67}$$

which unfortunately doesn't yield a closed-form expression. To gain further insight, we approximate the cumulative Gaussian function by the logistic sigmoid $\Phi(z) \approx \left(1+\exp\left(-C_\sigma z\right)\right)^{-1}$ with $C_\sigma = \pi^2/6$ to have matching slope at $z = 0$. After some algebra, this results in

$$C_2(P^+,t) \approx \sqrt{\sigma_\mu^{-2}+t}\frac{1}{C_\sigma}\log\frac{P^+}{1-P^+}, \tag{68}$$

showing that it becomes insufficient to use a shift of the accumulation starting point, as for the previous prior. Instead, we require both a shifted starting point, as well as an additional shift in the accumulated evidence that varies over time.

#### The relation between decision confidence and choice accuracy for biased priors

For either choice of the prior, the solutions that regains the boundary - decision correspondence result in a decision confidence that is the same at both boundaries, as long as these boundaries are symmetric around zero. For example, for the posterior Eq. (62) this is easy to see by replacing $\sigma_\mu^{-2}m + z$ by $\tilde{z}$, and, at the decision by $\tilde{z} = \pm\theta(t)$, depending on which choice has been made. This is seemingly at odds with expecting a different choice accuracy at either boundary, imposed by the biased prior.

To show that no overall inconsistency between choice accuracy and choice confidence exists, let us consider the simpler case of a prior with a single "difficulty" $\mu_0$, which is given by

$$p(\mu) = \frac{P^+}{2}\delta(\mu - \mu_0) + \frac{1-P^+}{2}\delta(\mu + \mu_0), \tag{69}$$

where $\delta(\cdot)$ is the Dirac delta function. That is, $\mu = \mu_0$ with probability $P^+$, and $\mu = -\mu_0$ with probability $1 - P^+$. With this prior, it is easy to show that the posterior becomes

$$p(\mu = \mu_0|z,t) = p(\mu \geq 0|z,t) = \frac{1}{1 + e^{-2\mu_0\left(z - \frac{1}{2\mu_0}\log\frac{P^+}{1-P^+}\right)}}. \tag{70}$$

For symmetric boundaries at $\pm\theta$, rather than shifting the accumulation starting point, we can equivalently shift the boundaries by the same amount to

$$\theta^+ = \theta - \frac{1}{2\mu_0}\log\frac{P^+}{1-P^+}, \qquad \theta^- = -\theta - \frac{1}{2\mu_0}\log\frac{P^+}{1-P^+}, \tag{71}$$

again leading to a constant decision confidence $\left(1 + \exp\left(-2\mu_0\theta\right)\right)^{-1}$ at either boundary.

To show that this decision confidence equals the probability of making the correct choice on average, we find this probability for each possible latent state value, using known expression for boundary hitting probabilities for diffusion models with asymmetric boundaries, as given in [4, 13]. For $\mu = \mu_0$, the upper boundary $\theta^+$ leads to the correct choice. This boundary is reached with probability

$$p\left(z = \theta^+ | z \in \left\{\theta^+, \theta^-\right\}, \mu = \mu_0\right) = \frac{e^{2\mu_0\theta} - \frac{1-P^+}{p^+}}{e^{2\mu_0\theta} - e^{-2\mu_0\theta}}, \tag{72}$$

which is the probability of making correct choices if $\mu = \mu_0$. Note that, unlike the confidence, this probability is modulated by $P^+$. In particular, it grows with an increase in $P^+$. In other words, the larger the a-priori probability that the upper boundary leads to the correct choice, the larger the probability that the decision maker chooses correctly in trials in which the upper boundary is indeed the correct choice.

For $\mu = -\mu_0$, the lower boundary $\theta^-$ leads to correct choices, which happens with probability

$$p\left(z = \theta^- | z \in \left\{\theta^+, \theta^-\right\}, \mu = -\mu_0\right) = \frac{e^{2\mu_0\theta} - \frac{P^+}{1-P^+}}{e^{2\mu_0\theta} - e^{-2\mu_0\theta}}, \tag{73}$$

where the only difference to the previous expression is the impact of the prior. Specifically, this probability shrinks with an increasing $P^+$.

The average probability of choosing correctly is a combination of both bound-hitting probabilities, weighted by the latent state probabilities, which, after some algebra, results in

$$p(\text{correct}) = p\left(z = \theta^+ | z \in \left\{\theta^+, \theta^-\right\}, \mu = \mu_0\right) p(\mu = \mu_0) + p\left(z = \theta^- | z \in \left\{\theta^+, \theta^-\right\}, \mu = -\mu_0\right) p(\mu = -\mu_0) = \frac{1}{1 + e^{-2\mu_0\theta}}, \tag{74}$$

where we have used $p(\mu = \mu_0) = P^+$ and $p(\mu = -\mu_0) = 1 - P^+$. This demonstrates that, even though the decision confidence differs from the probability of making the correct choices for individual choices, it equals the average probability of making correct choices. This unintuitive result follows from conditioning the choice probabilities on the latent state, which is unknown to the decision maker, and thus cannot be reflected in their decision confidence. Once this latent state is marginalized out (by averaging over it in Eq. (74)), consistency with the decision confidence is restored [7]. The same principle applies to the more complex priors used further above, but for those, it becomes hard to establish the equivalence between choice probability and decision confidence analytically.

#### Generating correlated momentary evidence

Recall that, for a given latent state $\mu$, the mutli-dimensional momentary evidence is drawn according to

$$\delta\boldsymbol{x}|\mu \sim \mathcal{N}\left(\left(\boldsymbol{a}\mu + \boldsymbol{b}\right)\delta t, \boldsymbol{\Sigma}\delta t\right), \tag{75}$$

where the parameters $\boldsymbol{a}$, $\boldsymbol{b}$ and $\boldsymbol{\Sigma}$ satisfy $\boldsymbol{a}^T\boldsymbol{w} = 1$, $\boldsymbol{b}^T\boldsymbol{w} = 0$, and $\boldsymbol{w}^T\boldsymbol{\Sigma}\boldsymbol{w} = 1$ (see Sec. "High-dimensional momentary evidence").

We satisfy the requirement on $\boldsymbol{a}$ and $\boldsymbol{b}$ by choosing

$$\boldsymbol{a} = \frac{\boldsymbol{w}}{\boldsymbol{w}^T\boldsymbol{w}}, \qquad \boldsymbol{b} = f_0\left(\boldsymbol{1} - \frac{\boldsymbol{1}^T\boldsymbol{w}}{\boldsymbol{w}^T\boldsymbol{w}}\boldsymbol{w}\right), \tag{76}$$

where $f_0$ is a parameter. The expression for $\boldsymbol{b}$ minimizes $\|\boldsymbol{b} - f_0\mathbf{1}\|$ under the constraint $\boldsymbol{b}^T\boldsymbol{w} = 0$, effectively introducing an approximate baseline at $f_0$.

For our choice for the covariance we were guided by observations that the noise covariance spectrum in neural population recordings has few dominant components, and otherwise rapidly drops towards small values. We achieve this while satisfying $\boldsymbol{w}^T\boldsymbol{\Sigma}\boldsymbol{w} = 1$ by designing a $\boldsymbol{\Sigma}$ that has one eigenvector $\boldsymbol{w}/\|\boldsymbol{w}\|$ with associated eigenvalue $1/\boldsymbol{w}^T\boldsymbol{w}$, and otherwise the desired eigenspectrum. To do so, we fill a $J \times J$ matrix $\boldsymbol{B}$ ($J$ is the size of $\delta\boldsymbol{x}$) with zero mean unit variance Gaussian random numbers, except for the first row, which we set to $\boldsymbol{w}$. This is followed by Gram-Schmidt orthonormalization of $\boldsymbol{B}$, such that the first row becomes $\boldsymbol{w}/\|\boldsymbol{w}\|$, while all other rows unit vectors, orthogonal to $\boldsymbol{w}$. We then choose a diagonal $\boldsymbol{D}$ with the first diagonal element $d_{11} = 1/\boldsymbol{w}^T\boldsymbol{w}$, and all other diagonal elements $d_{jj} = \max\left\{\sigma_x^2 e^{-j+1}, \sigma_0^2\right\}/\boldsymbol{w}^T\boldsymbol{w}$, with parameters $\sigma_x^2$ and $\sigma_0^2$. The final covariance matrix is then given by $\boldsymbol{\Sigma} = \boldsymbol{B}\boldsymbol{D}\boldsymbol{B}^T$.

If the weights change across consecutive trials $n-1$ and $n$, the momentary evidence needs to satisfy $\boldsymbol{a}_n^T\boldsymbol{w}_n = 1$, $\boldsymbol{b}_n^T\boldsymbol{w}_n = 0$, and $\boldsymbol{w}_n^T\boldsymbol{\Sigma}_n\boldsymbol{w}_n$ in each trial. For $\boldsymbol{a}_n$ and $\boldsymbol{b}_n$ this is easily achieved by re-computing them in each trial according to the above expressions.

The generation of $\boldsymbol{\Sigma}_n$ relies on a stochastic process, such that re-generating a new $\boldsymbol{\Sigma}_n$ in each trial might lead $\boldsymbol{\Sigma}_\cdot$ to change significantly across trials despite only small changes in $\boldsymbol{w}_\cdot$. To avoid this, we instead modify $\boldsymbol{\Sigma}_{n-1}$ by finding the smallest rotation $\boldsymbol{U}$ of $\boldsymbol{\Sigma}_{n-1}$ that satisfies $\boldsymbol{w}_n^T\boldsymbol{\Sigma}_n\boldsymbol{w}_n = 1$. To do so, we aim at finding $\boldsymbol{U}$ that satisfies $\boldsymbol{w}_n \propto \boldsymbol{U}\boldsymbol{w}_{n-1}$. This leaves $\boldsymbol{U}$ underconstraint. To introduce additional constraints, we would like to restrict the rotation imposed by $\boldsymbol{U}$ to the $(\boldsymbol{w}_{n-1}, \boldsymbol{w}_n)$ plane. We express this by using $\boldsymbol{\psi}_3, \dots, \boldsymbol{\psi}_J$ that are orthonormal unit vectors that are also orthogonal to $\boldsymbol{w}_{n-1}$ and $\boldsymbol{w}_n$, which we can find by Gram-Schmidt orthonormalization. For those vectors, we desire $\boldsymbol{\psi}_n = \boldsymbol{U}\boldsymbol{\psi}_n$. Overall, this leads to the linear equation

$$\boldsymbol{U}\begin{pmatrix} \frac{\boldsymbol{w}_{n-1}}{\|\boldsymbol{w}_{n-1}\|} & \frac{\boldsymbol{w}_n}{\|\boldsymbol{w}_n\|} & \boldsymbol{\psi}_3 & \dots & \boldsymbol{\psi}_J \end{pmatrix} = \begin{pmatrix} \frac{\boldsymbol{w}_n}{\|\boldsymbol{w}_n\|} & \frac{\tilde{\boldsymbol{w}}}{\|\tilde{\boldsymbol{w}}\|} & \boldsymbol{\psi}_3 & \dots & \boldsymbol{\psi}_J \end{pmatrix}, \tag{77}$$

where $\tilde{\boldsymbol{w}}$ is given by

$$\tilde{\boldsymbol{w}} = 2\frac{\boldsymbol{w}_{n-1}^T\boldsymbol{w}_n}{\boldsymbol{w}_n^T\boldsymbol{w}_n}\boldsymbol{w}_n - \boldsymbol{w}_{n-1}, \tag{78}$$

and which we can easily solve for[2] $\boldsymbol{U}$. With this rotation matrix, $\boldsymbol{\Sigma}_n$ is given by

$$\boldsymbol{\Sigma}_n = \frac{\boldsymbol{w}_{n-1}^T\boldsymbol{w}_{n-1}}{\boldsymbol{w}_n^T\boldsymbol{w}_n}\boldsymbol{U}\boldsymbol{\Sigma}_{n-1}\boldsymbol{U}^T, \tag{79}$$

where the re-scaling by the fraction ensures the correct scaling of the eigenvalues.

#### Simulation details

We used parameters $\sigma_0^2 = 0.001$, $\sigma_x^2 = 2$ and $f_0 = 0$ to generate the momentary evidence $\delta\boldsymbol{x}|\mu$, as described in the previous section. At the beginning of each trial sequence we drew the true weights according to $\boldsymbol{w} \sim \mathcal{N}(\boldsymbol{m}_w, \boldsymbol{S}_w)$, with unit mean $\boldsymbol{m}_w = \mathbf{1}$ and identity covariance $\boldsymbol{S}_w = \boldsymbol{I}$. For that sequence, the diffusion model bounds $\pm\theta$ were time-invariant, and tuned to maximize the reward rate if the true weights were used to combined the inputs. We used $\sigma_\mu^2 = 3^2$ to draw $\mu$ in each trial. This $\mu$ determined the correct choice by $y^* = 1$ if $\mu \geq 0$, and $y^* = -1$ otherwise. The reward rate was given by

$$RR = \frac{p(\text{correct}) - c_{accum}\langle t\rangle}{\langle t\rangle + t_{iti}}, \tag{80}$$

where the average was across trials, and we set the evidence accumulation cost to $c_{accum} = 0.01$ and the inter-trial interval to $t_{iti} = 2s$. For non-stationary weights, we re-drew the weights after each choice according to Eq. (46), with $\boldsymbol{A} = \lambda\boldsymbol{I}$, $\boldsymbol{b} = (1-\lambda)\boldsymbol{m}_w$, and $\boldsymbol{\Sigma}_d = \left(1-\lambda^2\right)\boldsymbol{S}_w$, and set the decay factor to $\lambda = 1 - 0.01$. This yields a weight diffusion that follows a first-order autoregressive process with steady-state mean $\boldsymbol{m}_w$ and covariance $\boldsymbol{S}_m$.

To compare the weight learning performance of ADF to alternative models, we simulated 1,000 learning trials 5,000 times, and reported the reward rate per trial averaged across these 5,000 repetitions. To assess steady-state performance, we performed the same procedure with non-stationary weights, and reported reward rate averaged over the last 100 trials, and over 5,000 repetitions. The sequential choice dependencies in Fig. 4a/b where also computed from these last 100 trials. The learning rate in Fig. 1d in the main text shows the pre-factor to $\boldsymbol{\Sigma}_w\tilde{\boldsymbol{x}}$ in Eq. (41) over decision confidence for a subsample of the last 10,000 trials of a single 15,000 trial simulation with non-stationary weights. For the Gibbs sampler, we drew 10 burn-in samples, followed by 200 samples in each trial. For the particle filter we simulated 1,000 particles.

We sped up the diffusion model simulations by simulating the diffusion directly in the one-dimensional $\boldsymbol{w}^T\boldsymbol{x}_n(t)$ space. This resulted in a one-dimensional diffusion model whose first-passage time distribution is known and can be efficiently drawn from [5]. The final $\boldsymbol{x}_n(t_n)$ was recovered by drawing it from

$$x_n(t_n) \sim \mathcal{N}\left(\frac{\mu_n t_n}{\boldsymbol{w}^{*T}\boldsymbol{w}^*}\boldsymbol{w}^*, \frac{t_n}{\boldsymbol{w}^{*T}\boldsymbol{w}^*}\boldsymbol{I}\right), \tag{81}$$

subject to the constraint $\boldsymbol{w}^T\boldsymbol{x}_n(t_n) = y_n\theta$, and where $\boldsymbol{w}^*$ and $\boldsymbol{w}$ denote the true weights, and the weights used for evidence accumulation, respectively.

[2] Most likely there exists a closed-form expression for $\boldsymbol{U}$. We found it by solving the above expression numerically in each trial.

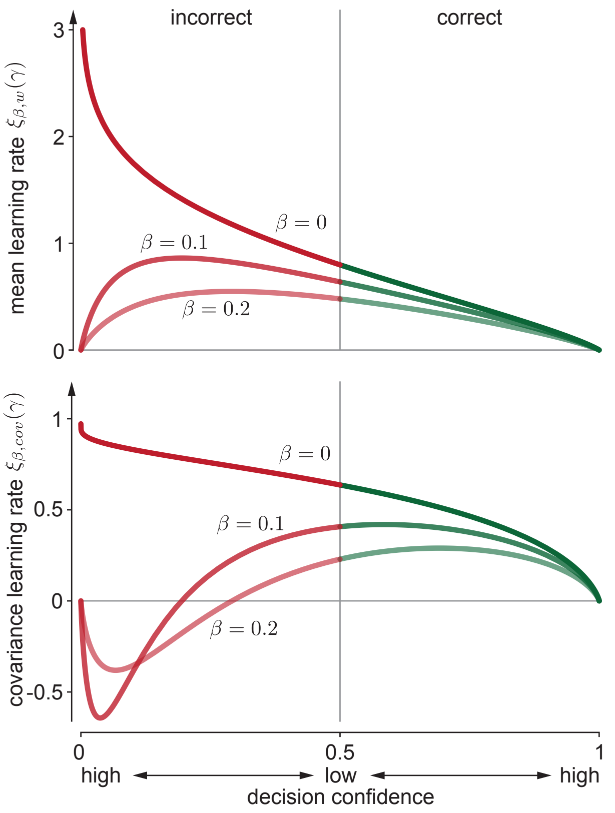


Figure 1: Assumed density filtering learning rate modulators for noise-free and noisy feedback $y^*$. The top panel shows the learning rate modulator $\xi_{\beta,w}(\gamma)$ of the mean update for different levels of feedback noise, $\beta$. The bottom panel shows the same for the learning rate modulator $\xi_{\beta,cov}(\gamma)$ of the covariance update. In both cases, the marginal decision confidence associated with the feedback $p(y^*|\boldsymbol{x}, t) = \Phi(\gamma)$ is varied along the horizontal axis. This marginal decision confidence is $> 1/2$ for correct (green), and $< 1/2$ for incorrect (red) choices. $\beta = 0$ corresponds to the noise-free case, for which $\xi_w(\gamma) = \xi_{\beta,w}(\gamma)$ and $\xi_{cov}(\gamma) = \xi_{\beta,cov}(\gamma)$.

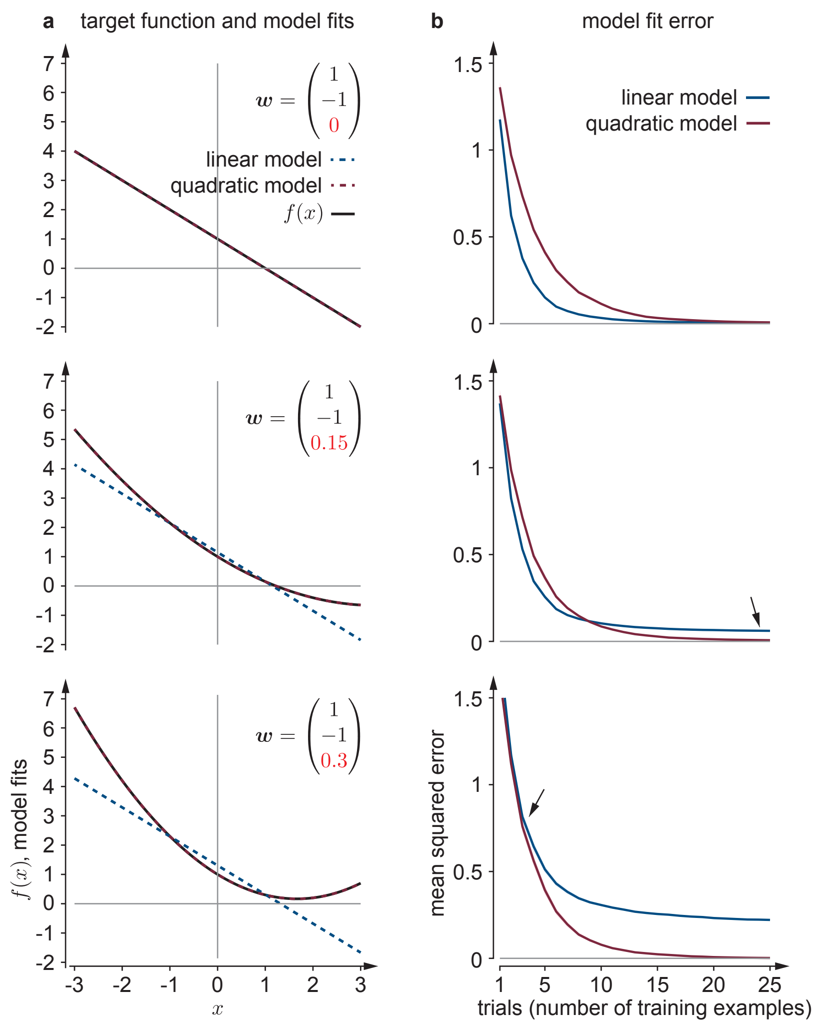


Figure 2: Simpler models can learn more rapidly than more complex models, even if they are unable to provide perfect fits. We show model fits and model error for a linear (blue) and quadratic (red) model when fitting the target function $f(x) = w_1 + w_2 x + w_3 x^2$ for different $w_3$'s (red number in the $\boldsymbol{w}$'s in (a)) for the top, middle, and bottom row. The quadratic model has the same functional form as $f(x)$ and learns all $\boldsymbol{w}$. The linear model fixes $w_3 = 0$, and only learns $w_1$ and $w_2$. Both models are fitted to training data consisting of $(x_n, f(x_n))$-pairs, by finding the model weights that minimize the mean squared error between model predictions and given $f(x_n)$'s across all observed $x_n$'s. (a) With $10^4$ training examples, both the linear and the quadratic model can fit a linear function (top; model fits and target function plotted on top of each other). As soon as the target function becomes quadratic (middle & bottom), the linear model fails to perfectly fit this function. (b) The mean squared error, here shown as an average across 500 repetitions across different training sets, drops more rapidly for the linear model than for the quadratic model if the target function is linear (top). This is because the linear model needs to learn fewer parameters for the same training set size. The error of both models goes to zero once the training set size increases. Even if the target function becomes quadratic (middle), the linear model can still learn more rapidly than the quadratic model (blue initially drops faster than red), even if it can't reduce its error to zero (arrow). This is only possible if the target function is still close-to-linear over the range of interest. Once it becomes too non-linear (bottom), the linear model learns slower than the quadratic model (arrow), and features a significantly worse asymptotic error. In (b), the mean squared error was in each repetition and for each training set size computed over 1000 new $x$'s that were not part of the training set. For all simulations, the $x$'s were drawn from $x \sim \mathcal{N}(0, 1)$. All learning was performed through optimization, by minimizing the mean squared error. We could have equally used learning by inference (using Bayesian linear regression with sufficiently uninformative priors), without affecting the results. Therefore, the shown effects are independent of the chosen learning formalism.
